# TwinSAR: An Adaptive Kernel-based Algorithm with logit-transformed Z-score Filtering for Chemical Twin Detection in Large-scale Virtual Screening

**DOI:** 10.64898/2026.05.12.724687

**Authors:** Haris Kulosmanović, Cem Uğuz, Serdar Durdağı

## Abstract

Molecular similarity searching is a workhorse of cheminformatics, but the dominant Tanimoto/topological-fingerprint paradigm has well-known blind spots. It is highly sensitive to molecular size, suffers from steep activity cliffs, and frequently fails to retrieve scaffold-hopping bioisosteres. A complementary descriptor that has received comparatively little attention is global elemental composition. Despite the conceptual simplicity of comparing molecules by their elemental ratios, no widely deployed method exists for the statistically rigorous identification of “chemical twins” defined by stoichiometric proximity. We address this gap with TwinSAR (Stoichiometric Analysis and Retrieval), an adaptive kernel-based algorithm that combines three methodological innovations: (i) binary fingerprint blocking that partitions molecule by element-presence patterns and bounds the cost of all-pairs comparison from O(*NM*) to *O*(∑*n*_*i*_*m*_*i*_) enabling million/billion-scale searches; (ii) a per-block adaptive radial basis function (RBF) kernel whose precision parameter is calibrated independently for each fingerprint block via the median heuristic, providing fair similarity comparison across chemical sub-spaces of vastly different density; and (iii) a logit-transformed Z-score filter that maps bounded RBF scores onto an unbounded scale, allowing high-similarity pairs to be prioritized relative to the empirical score distribution of their own fingerprint block. TwinSAR is offered in two operating modes: (i) a deterministic BULK mode for exact reproducibility; and (ii) a stochastic FAST mode that achieved a 3.29× wall-clock speed-up in the present benchmark while preserving the similar unique-query and unique-target coverage. Statistical validation showed that detected twin pairs are 12.7× more similar in absolute ratio space than block-matched random pairs (*p* < 0.001), while a column-permutation negative control returned a median of zero spurious twins across three independent permutations. A controlled benchmark further established that an 8-element representation (single-element heavy-atom ratios) is sensitivity-equivalent to a comprehensive 254-element representation while running 3.55× faster. As a case study, TwinSAR was deployed in an end-to-end virtual screening pipeline against the BCL-2 target protein, where it reduced a 327,071-compound commercial library to a 390 focused candidate panel. The chemical interpretability of the retrieved twins is illustrated by their structural diversity around conserved heavy-atom skeletons. TwinSAR therefore provides a fast, conformation-free, and statistically principled prefilter that is fully orthogonal to topological fingerprints.

## 1. Introduction

Molecular similarity is a central organizing principle in cheminformatics and ligand-based drug discovery.^1–3^ The practical assumption that structurally or physicochemically similar molecules may display related biological activities has motivated the development of similarity searching, clustering, virtual screening, and structure–activity relationships (SAR) modeling strategies across large chemical libraries.^4-6^ In contemporary workflows, two-dimensional (2D) topological fingerprints (i.e., Morgan/ECFP, MACCS, FCFP) combined with the Tanimoto coefficient remain the *de facto* standard for similarity assessment.^1–3^ These descriptors are inexpensive to compute and excel at retrieving close structural analogues, but they suffer well-documented limitations: (i) Tanimoto similarity is highly sensitive to molecular size, exhibits steep activity cliffs,^7^ and frequently fails to identify scaffold-hopping bioisosteres that share pharmacophoric features but differ in topology.^8,9^ (ii) Pharmacophore- and shape-based methods such as ROCS^10^ partially address scaffold hopping but require three-dimensional (3D) conformer generation that is computationally expensive at the scale of millions of molecules.

A complementary, lower-dimensional descriptor that has received comparatively little attention is the elemental composition of the molecule. Two compounds can share a near-identical heavy-atom stoichiometry while differing markedly in connectivity, and conversely, highly potent inhibitors of a given target frequently cluster in narrow regions of stoichiometric space because productive interactions with a fixed binding site impose constraints on hydrogen-bond donor/acceptor counts, aromaticity, and the inclusion of specific halogens.^11,12^ Stoichiometric similarity thus offers a fast, conformation-free, scaffold-agnostic filter that is orthogonal to fingerprint-based metrics. Despite this conceptual simplicity, to the best of our knowledge, no widely deployed method exists for the statistically rigorous identification of “chemical twins” defined by elemental ratios. The challenges that any such method must overcome are three-fold: (i) bounding the cost of all-pairs comparison over libraries that already exceed 10^9^ compounds; (ii) scoring similarities adaptively despite very heterogeneous distance distributions across different chemical sub-spaces, ranging from tightly clustered classes such as simple aromatic carboxylic acids to broadly distributed halogenated heterocycles; and (iii) distinguishing exceptional high-similarity pairs from incidental matches by ranking each pair relative to the empirical score distribution of its own stoichiometric block.

In this study, we introduce TwinSAR (Stoichiometric Analysis and Retrieval), an adaptive kernel-based algorithm that addresses these three challenges through three corresponding methodological innovations. First, binary fingerprint blocking partitions molecules by their element-presence pattern and restricts comparisons to within-block pairs, reducing the expected complexity from O(*NM*) to *O*(∑*n*_*i*_*m*_*i*_) and yielding one to two orders of magnitude in speed-up on heterogeneous libraries. Second, a per-block adaptive RBF kernel calibrates its precision parameter *γ* independently for each fingerprint block using the median heuristic,^13^ enabling fair similarity comparison across chemical sub-spaces of very different density. Third, a logit-transformed Z-score filter that maps bounded RBF scores onto an unbounded scale and provides a block-normalized ranking statistic for identifying pairs that lie in the extreme high-similarity tail of their local stoichiometric background.^14^ In the present benchmark, FAST mode reproduced the BULK mode molecule-level coverage while providing a substantial speed-up. The algorithm is validated by three orthogonal statistical tests and demonstrated in a virtual screening application against the BCL-2 target protein, where it reduces a 327,071-compound SPECS small molecule library to a focused candidate panel. The structural diversity of the retrieved twins around conserved heavy-atom skeletons supports the chemical interpretability of the algorithm.

## 2. Methods

### 2.1. The TwinSAR algorithm

#### 2.1.1. Stoichiometric feature generation

Each molecule M is represented by a vector of heavy-atom element ratios over the non-hydrogen atom set Ω = {C, N, O, F, S, Br, Cl, I}, deliberately omitting hydrogen because of its sensitivity to protonation and tautomeric assignment.^14^ The streamlined 8-element representation (*single8*) defines ratio_X_ = count(*X*) / *N*_heavy_ for *X* ∈ Ω, where *N*_heavy_ denotes the total heavy-atom (i.e., non-hydrogen) count of M, yielding exactly eight features per molecule. A more comprehensive 254-element representation (*full254*) encodes all non-trivial subsets of Ω, producing 2^8^ − 2 = 254 features (excluding the empty set and the full set). Because, higher-order subset ratios are deterministic functions of the eight single-element ratios, the *full254* representation was evaluated primarily as a stress test of whether redundant higher-order stoichiometric features alter retrieval behavior. The empirical comparison of the two representations is reported in Section 3.3.

#### 2.1.2. Binary fingerprint blocking

Naive all-pairs comparison of *N* query molecules against *M* targets has time complexity O(*NM*), which becomes prohibitive at the scale of 10^9^ compounds. To bound this cost, we adopt a binary fingerprint blocking strategy: for every molecule, a fingerprint b ∈ {0,1}^d^ is constructed by setting *b*_i_ = 1 if and only if the *i*-th ratio feature is non-zero (*d* = 8 for *single8, d* = 254 for *full254*). Two molecules are eligible for direct comparison only if their fingerprints are identical, equivalent to requiring exact agreement on the set of elements present. This partitions the dataset into disjoint blocks indexed by fingerprint, reducing the expected complexity to *O*(∑*n*_*i*_*m*_*i*_), where *n*_i_ and *m*_i_ are the number of queries and targets in block *i*. Empirically these yield one to two orders of magnitude in speed-up on heterogeneous chemical libraries while preserving completeness for stoichiometrically meaningful pairs. The blocking strategy also has the desirable property of guaranteeing that no comparison is performed between molecules with disjoint elemental compositions, which would in any case produce uninformative similarities.

#### 2.1.3. Per-block adaptive RBF kernel

Within each block, similarity between feature vectors *x* and *y* is computed using the Gaussian radial basis function (RBF) kernel *K*(*x, y*) = *exp*(−γ ∥ *x* − *y* ∥^”^). Critically, the kernel precision parameter *γ* is not a global hyperparameter but is calibrated per block via the median heuristic^13^: *γ* = 1/median(‖x−y‖^2^+ε), where ε is a small constant added for numerical stability to prevent division-by-zero errors in degenerate cases. This produces an interpretable scaling: two molecules separated by the median squared distance always receive a similarity of *e*^−1^ ≈ 0.368 regardless of the absolute spread of the block. The chemical heterogeneity of fingerprint blocks varies enormously, from tightly clustered classes such as simple aromatic carboxylic acids to broadly distributed halogenated heterocycles, and per-block adaptation is essential for fair similarity comparison; using a single global *γ* would either saturate dense blocks at similarity ≈ 1 or render sparse blocks effectively flat at similarity ≈ 0.

#### 2.1.4. Logit transformation and Z-score filtering

Raw RBF similarities are bounded in [0, 1] and are heavily compressed near 1 for highly similar pairs, producing pronounced skew and bounded-score compression that make direct global thresholding difficult. To map similarities into an unbounded space amenable to *Z*-score thresholding, we apply the logit transform *L*(*s*) = ln[*s*/(1−*s*)],^14^ where *s* denotes the raw similarity; to ensure numerical stability, similarity values of s=1 are clipped to 1-10^−7^ to prevent singularity. Within each fingerprint block, the logit-transformed values were standardized as *Z(s) = [L(s) − μL] / σL*, where *μL* and *σL* are the mean and standard deviation of *L(s)* in that block. Candidate twin pairs were retained using a stringent threshold of *Z* > 2.576, which nominally corresponds to the upper 0.5% tail under a standard normal approximation; empirical per-block calibration indicates that this threshold is conservative in the majority of blocks (empirical false-positive rate below the nominal 0.5% in most blocks), although a minority of smaller or high-kurtosis blocks exceed this nominal rate, as detailed in the Table S1. The impact of the logit transformation on the score distribution is illustrated in Figures S1 and S2.1, where the raw similarity distribution, highly compressed near 1.0, is contrasted with the logit-transformed Z-score distribution. This transformation effectively reduces bounded-score compression and enables statistically rigorous, block-wise tail prioritization. The TwinSAR framework was further validated through a series of technical assessments, including kernel sensitivity analysis (Figure S2.1), comprehensive regression model diagnostics and feature importance (Figures S2.2 and S5), and quantitative enrichment benchmarking (Figure S2.3). Furthermore, the computational efficiency of the blocking strategy was evaluated (Figure S2.4), with full structural and binding results for the BCL-2 case study detailed in the supplemental results (see Figure S2.5 and see Tables S4–S8). Collectively, these analyses demonstrate that TwinSAR provides a robust and computationally efficient platform for identifying bioactive stoichiometric twins across diverse chemical spaces. This block-wise standardization ensures that retained twin pairs are statistically more similar than the typical pair within their own stoichiometric class. Because pairwise comparisons are not independent and no explicit multiple-testing correction is applied, this threshold should be interpreted as a stringent empirical tail-selection heuristic rather than as a formal multiple-testing-adjusted false-discovery-rate procedure. The empirical validity of this heuristic is supported by the column-permutation negative control (Section 2.2), which returned a median of zero spurious twins across three independent permutations and a worst-case spurious-pair rate of 0.94% of detected BULK pairs (Section 3.4), providing a dataset-specific empirical upper bound on the false-positive rate in lieu of a formal correction. The effect of the logit transformation on the score distribution is illustrated in Figure S1, where the raw similarity distribution, highly compressed near 1.0, is contrasted with the logit-transformed *Z*-score distribution, which reduces bounded-score compression and enables block-wise tail-based prioritization.

#### 2.1.5. Operating modes: BULK vs. FAST

TwinSAR provides two operating modes that differ exclusively in the calculation of *γ*. In BULK mode (deterministic), *γ* is computed using the squared distances of all intra-block pairs, guaranteeing byte-identical results across runs. In FAST mode (non-deterministic), *γ* is estimated from a uniform random sample of at most *MAX_GAMMA_SAMPLES* = 5,000 squared distances per block. As characterized quantitatively in Section 3.4, the two modes showed highly similar statistical behavior in the present benchmark, with FAST providing a substantial speed-up at the cost of minor stochastic variability; recommended use cases are summarized in Table S3.

#### 2.1.6. Twin detection workflow

Given a query set *Q* and a target set *T*, the TwinSAR workflow proceeds as follows: (1) compute ratio features and fingerprints for *Q* ∪ *T*; (2) partition *Q* and *T* by fingerprint block; (3) for each block in which both *Q* and *T* have at least one molecule, calibrate *γ* (BULK or FAST) and compute the RBF similarity for every (*q, t*) pair; (4) apply the logit + *Z*-score filter; and (5) return the surviving (*q, t*) pairs together with their similarity, *Z*-score, and block identifier.

### 2.2. Statistical validation framework

Algorithmic validity was established with three orthogonal tests. First, the twin versus random ratio similarity test compares the mean absolute ratio difference between detected twin pairs and random pairs drawn from the same fingerprint block; a robust algorithm should produce twins that are significantly more similar than random pairs. Second, the column-permutation negative control applies a random permutation π of the eight ratio columns uniformly to every target molecule, so that values originally stored in the “ratio C” column are reassigned to, e.g., “ratio I”, those in “ratio N” to “ratio S”, and so on. This procedure disrupts the chemical correspondence between element identities and their ratio values; importantly, while the multivariate composition of the target library is preserved across molecules, each column’s marginal distribution is reassigned to a different element position rather than preserved in place, so the permutation does not constitute a row-wise within-column shuffle. The resulting element-label scrambling collapses the query–target block overlap from 17 shared blocks to between 1 and 13 surviving common blocks across permutations, reflecting the sensitivity of the binary fingerprint to element-identity reassignment. Three independent permutations were performed and the median number of retained twins reported. Third, the similarity–Z-score correlation measures whether the logit-transformed *Z*-score provides additional discriminative power beyond raw similarity; a moderate (rather than near-unity) correlation supports that block-wise normalization materially re-ranks pairs relative to their raw similarity, demonstrating the intended effect of the logit-Z standardization step.

### 2.3. Case study: BCL-2 inhibitor retrieval

To represent the utility of TwinSAR in a realistic virtual screening campaign, we deployed it as the primary retrieval step of an end-to-end pipeline against the anti-apoptotic protein B-cell lymphoma 2 (BCL-2).^15-17^ The reference query set was the BCL-2 activity dataset extracted from ChEMBL release 33, (https://www.ebi.ac.uk/chembl/) comprising 1,067 small molecules with experimentally measured IC_50_ values; for the production pipeline this set was pre-filtered to retain only high-activity binders (pIC_50_ > 6.5; *n* = 692). The screening library was the SPECS commercial collection (327,071 compounds, www.specs.net). SMILES strings were standardized, neutralized, and stripped of salts using RDKit^18^ prior to feature generation. After TwinSAR retrieval, candidates were filtered for drug-likeness using a relaxed Lipinski-derived rule set (MW ≤ 1000 Da; −2 ≤ logP ≤ 10; HBD ≤ 6; HBA ≤ 15; TPSA ≤ 250 Å^2^; rotatable bonds ≤ 20),^19,20^ ranked by predicted potency using a CatBoost^21^ pIC_50_ regression model, trained on the ChEMBL-BCL2 dataset using an [80/10/10 training/validation/test-set split], with hyperparameters selected by three-fold cross-validation on the training partition and performance evaluated on a held-out test set (test R^2^ = 0.848, RMSE = 0.738; validation–test gap ΔR^2^ = 0.011) indicated negligible overfitting, with feature importance analysis highlighting the dominance of topological connectivity and molecular weight in potency prediction (Figure S5; full diagnostics in Section S2). Finally evaluated by Boltz-2^22^ structure-based binding probability prediction against full-length and truncated BCL-2 receptor variants (FASTA sequences in Section S1, Supporting Information). The detailed results are reported in Section S3 (Supporting Information).

## 3. Results

### 3.1. Algorithm overview and implementation

The end-to-end TwinSAR algorithm and its embedding in a full virtual screening workflow are summarized in Figure 1. Phases 1 and 2 of the figures depicts the algorithmic core: feature generation and binary fingerprint blocking, followed by similarity scoring with the per-block adaptive RBF kernel and the logit-transformed *Z*-score statistical filter. Phase 3 illustrates one particular deployment of the algorithm, a BCL-2 inhibitor discovery funnel, used here to demonstrate utility on a real chemical library; the algorithm itself is target- and pipeline-agnostic. The TwinSAR algorithm was implemented in Python 3.11 using NumPy and SciPy for linear algebra, scikit-learn^23^ for variance thresholding and scaling, and RDKit^18^ for cheminformatics.

**Figure 1.**
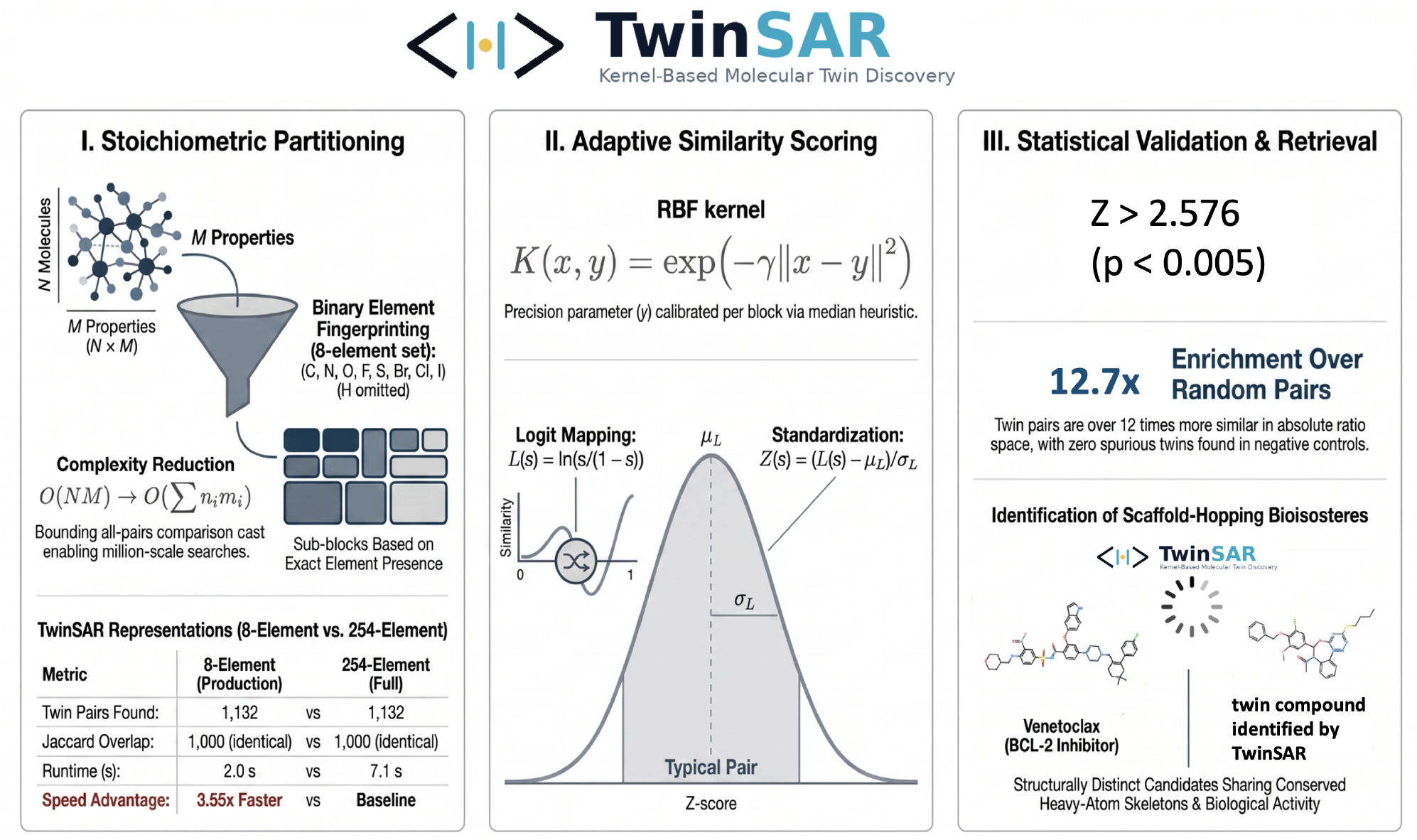
Overview of the TwinSAR algorithm and its embedding in a complete virtual screening workflow. Phase 1, each molecule is encoded as an *8-element* heavy-atom ratio vector and partitioned into a binary fingerprint block determined by which elements are present, bounding the cost of all-pairs comparison. Phase 2, within each block, similarity is scored with a Gaussian RBF kernel whose precision parameter γ is calibrated by the median heuristic, then bounded similarities are mapped onto an unbounded logit-Z scale, enabling block-wise prioritization of pairs in the extreme high-similarity tail of their local stoichiometric background. Phase 3, an end-to-end BCL-2 inhibitor discovery funnel that integrates TwinSAR retrieval with drug-likeness filtering, CatBoost pIC_50_ prediction, and Boltz-2 structure-based binding validation.

The source code is available from our research group’s github page: https://github.com/DurdagiLab/Twinsar

### 3.2. Statistical validation

The three orthogonal statistical tests defined in Section 2.2 were applied to TwinSAR (BULK mode, 8-element representation) on a query set of 692 high-activity ChEMBL-BCL2 molecules searched against the SPECS library; results are summarized in Table 1 and visualized in Figure 2.

**Table 1.** TwinSAR core validation metrics on the production query set.^*a*^.

| Metric | Result | Interpretation |
| --- | --- | --- |
| Twin vs. random ratio difference | 12.7× more similar | Pass |
| Negative control (column permutation) | 0 shuffled twins (median) | Pass |
| Similarity–Z-score correlation, BULK<sup>b</sup> | $r = 0.449$ | Moderate<sup>d</sup> |
| Similarity–Z-score correlation, FAST<sup>c</sup> | $r = 0.432$ | Moderate<sup>d</sup> |

<sup>a</sup> Production query set, $n = 692$ , pre-filtered to $\text{pIC}_{50} > 6.5$ ; 8-element representation; SPECS target library, 327 071 molecules; Pearson $r$ reported for the per-target similarity–Z-score consistency. <sup>b</sup> BULK uses the full block for $\gamma$ estimation. <sup>c</sup> FAST uses up to 5000 randomly sampled pairs per block. <sup>d</sup> Block-normalization effect confirmed.

**Figure 2.**
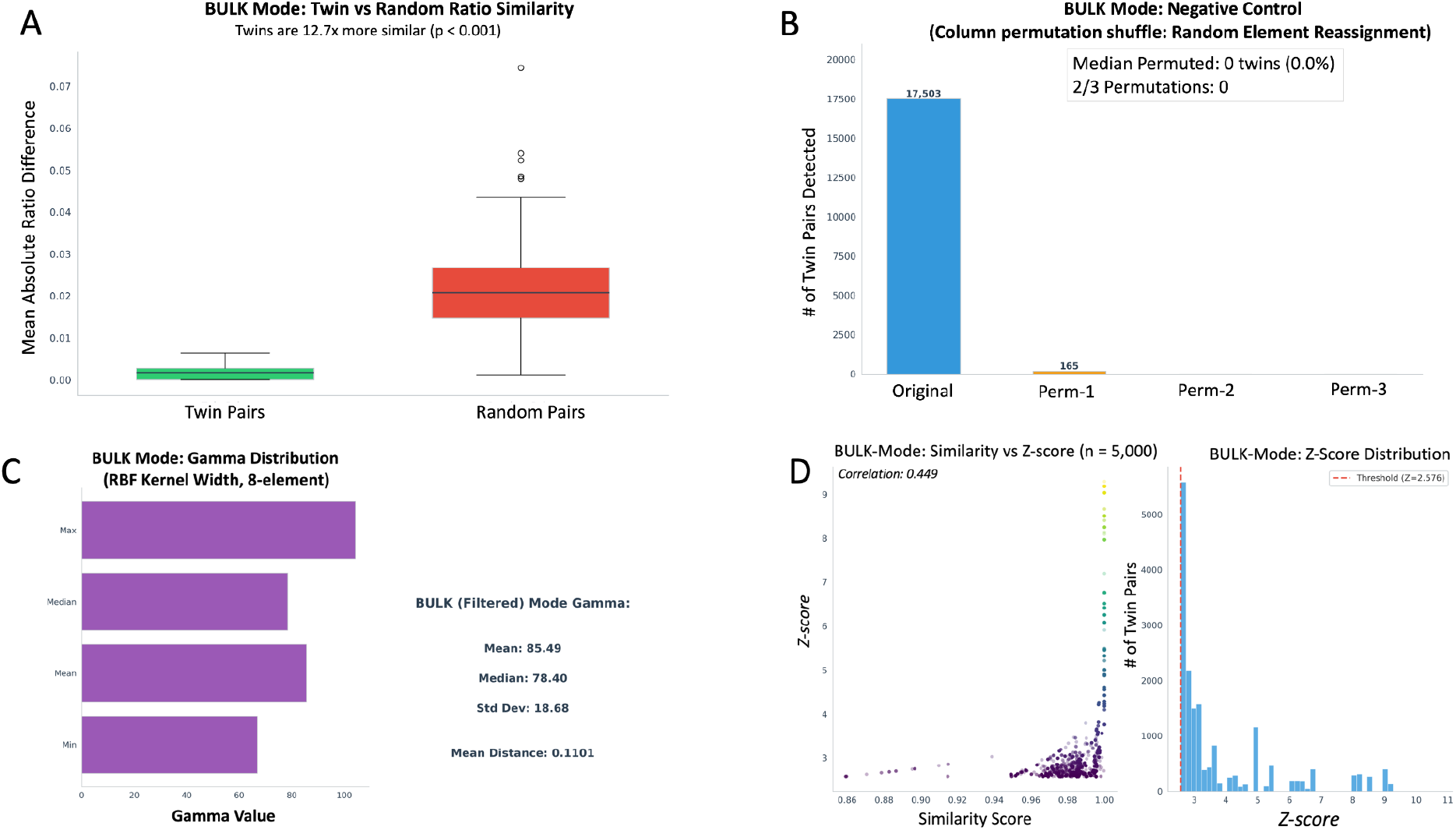
Statistical validation of TwinSAR (BULK mode, 8-element representation). (A) Twin pairs (green) are 12.7× more similar in ratio space than random pairs drawn from the same fingerprint block (red, p < 0.001). (B) Negative control via column-permutation shuffle: median permuted twins = 0, confirming the algorithm rejects scrambled element assignments. (C) Distribution of the adaptive kernel precision parameter γ across fingerprint blocks (mean = 85.49, median = 78.40); the broad block-to-block variation demonstrates the necessity of per-block calibration. (D) Per-target similarity vs. Z-score relationship (Pearson r = 0.449); the moderate correlation confirms that block-wise normalization materially re-ranks pairs relative to raw similarity, reflecting the intended effect of the logit-Z standardization step.

First, twin pairs detected against the SPECS library were 12.7× more similar in absolute ratio space than random pairs drawn from the same fingerprint block (p < 0.001; Figure 2a), confirming that the algorithm captures genuine stoichiometric proximity and not merely block-membership coincidence. Second, the column-permutation negative control yielded a median of 0 spurious twins across three independent permutations (Figure 2b; full results in Table S2). The small residual signal observed in Permutation 1 (165 pairs confined to a single surviving common block) reflects an accidental stoichiometric coincidence within that specific block under that particular element-label assignment, rather than a systematic signal. Critically, Permutations 2 and 3 produced zero spurious twins despite retaining 10 and 13 common blocks respectively, demonstrating that the number of surviving blocks does not predict spurious-twin yield; the residual in Permutation 1 is therefore attributable to the accidental structure of that one block rather than to any algorithmic artefact. The near-complete suppression across all three permutations (median = 0; worst-case 0.94% of detected BULK pairs) confirms that TwinSAR captures genuine stoichiometric proximity rather than coincidental block-level overlap. Third, the per-target similarity–*Z*-score correlation (*r* = 0.449 BULK mode; *r* = 0.432 FAST mode; Figure 2d) is moderate, confirming that block-wise normalization substantially re-ranks pairs relative to their raw similarity, consistent with the intended purpose of the logit-Z standardization step, as the Z-score reflects how exceptional a similarity is within its local block rather than an independent chemical descriptor. The Z-score acts as a precision filter, stripping away the broad background of incidental similarities and preserving only the extreme right-tail cases representing statistically exceptional pairs. The distribution of the adaptive kernel precision parameter *γ* across fingerprint blocks (Figure 2c; mean = 85.49, median = 78.40, range ∼30–110) confirms that the median heuristic spans an order-of-magnitude range across blocks, validating the necessity of per-block calibration: a single global γ cannot adapt to the heterogeneity of the chemical sub-spaces. Equivalent four-panel validation in FAST mode mirrors the BULK profile (Figure S3).

### 3.3. Representation robustness: 8-element vs. 254-element

A central design question for the algorithm is whether the streamlined 8-element ratio representation provides sufficient discriminative information, or whether the comprehensive 254-element subset-ratio representation captures additional categorical structure relevant to twin detection. We addressed this question with a controlled benchmark on 1,067 query molecules against 5,000 target molecules, running both representations under identical conditions; results are tabulated in Table 2 and visualized in Figure 3.

**Table 2.** Comparison of 8-element and 254-element TwinSAR representations on a controlled 1067 × 5000 benchmark.^*a*^.

| Metric | 8-element | 254-element |
| --- | --- | --- |
| Features used | 8 | 254 |
| Twin pairs found | 1132 | 1132 |
| Unique query molecules | 209 | 209 |
| Unique target twins | 512 | 512 |
| Mean similarity | 0.9916 | 0.9916 |
| Median similarity | 0.9959 | 0.9959 |
| Mean Z-score | 3.864 | 3.864 |
| Pairs with similarity > 0.99 | 835 | 835 |
| Runtime / s | 2.0 | 7.1 |
| Jaccard overlap (target twins) | 1.0000 | 1.0000 |

<sup>a</sup> Benchmark: 1067 query molecules screened against a 5000-molecule target library; Z-score and Jaccard overlap defined as in the main text. Identical results across all detection metrics (Jaccard = 1.000 for unique queries, target twins, and twin pairs) indicate that the 246 additional subset-ratio features of the 254-element representation provide no detection sensitivity beyond the eight single-element ratios in the chemical space tested.

**Figure 3.**
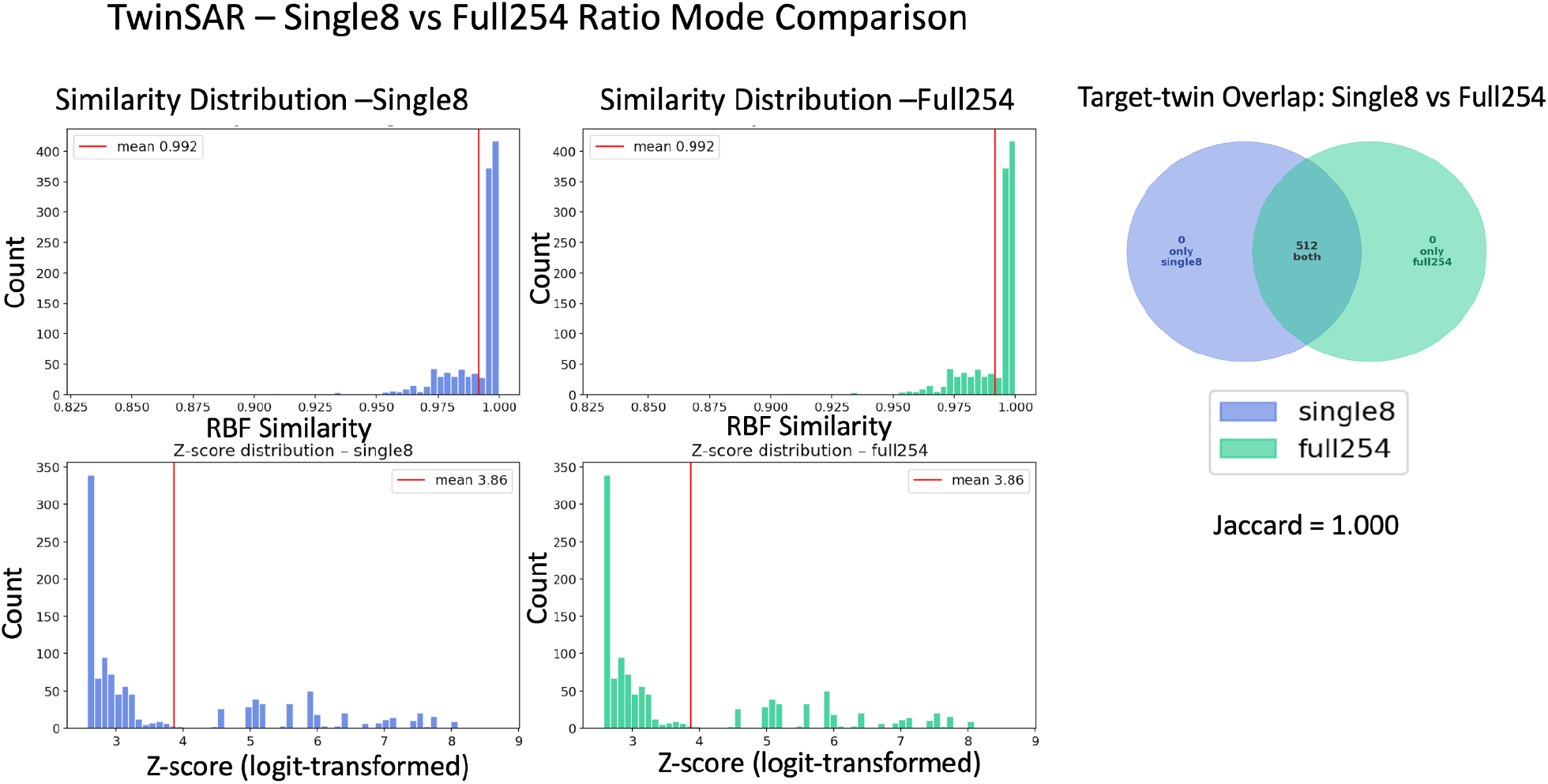
Comparison of 8-element (*single8*) and 254-element (*full254*) TwinSAR representations on the controlled benchmark. (Left, top to bottom) RBF similarity and *Z-score* distributions for *single8* (purple) and *full254* (green). (Right) Venn diagram of the unique target twins detected by each representation: all 512 are shared, with zero unique to either mode (Jaccard = 1.000). The 254-element representation incurred a 3.55× runtime penalty (7.1 s vs. 2.0 s) without retrieving a single additional pair, justifying the adoption of the streamlined 8-element representation as the production default.

Both representations produced exactly the same 1,132 twin pairs spanning the same 209 unique queries and 512 unique targets, with identical similarity distributions (mean = 0.9916), identical *Z*-score distributions (mean = 3.864), and a Jaccard overlap of 1.000 on the target-twin set. The 254-element representation incurred a 3.55× wall-clock penalty (7.1 s vs. 2.0 s) without retrieving a single additional pair. This result is consistent with the linear dependence of higher-order subset ratios on the eight single-element ratios in Euclidean geometry. Because subset-ratio features are derived from the same underlying elemental counts, their block-level information is largely redundant with the single-element presence fingerprint in the drug-like chemical space examined here. We therefore adopt *single-8* as the production default; *full-254* remains available for specialized applications where complex multi-element stoichiometric relationships may matter (e.g., organometallic libraries or coordination compounds).

### 3.4. Dual-mode characterization: BULK vs. FAST

The two operating modes were directly compared on the production set (692 queries × 327,071 targets) using the 8-element representation; results are summarized in Table 3. FAST mode achieved a 3.29× wall-clock speed-up over BULK (3.41 s vs. 11.24 s) while detecting all 17,503 BULK pairs plus 2,778 additional pairs. The two modes returned unique queries (BULK: 406, FAST: 420) and unique target twins (BULK: 10,387, FAST: 11,073), with FAST fully covering and expanding beyond BULK’s stoichiometric coverage of the chemical space.

**Table 3.** Dual-mode characterization of TwinSAR on the production set.^*a*^.

| Metric | BULK | FAST |
| --- | --- | --- |
| Processing time / s | 11.24 | 3.41 |
| Speed-up factor | 1.00× | 3.29× |
| Twin pairs | 17 503 | 20 281 |
| Unique query molecules | 406 | 420 |
| Unique target twins | 10 387 | 11 073 |
| BULK-only pairs | — | 0 |
| FAST-only pairs | — | 2778 |
| Jaccard overlap | 0.863 | 0.863 |

<sup>a</sup> Production set: 692 queries × 327 071 SPECS targets; 8-element representation; single-thread CPU. FAST achieves a 3.29× speed-up over BULK while fully containing the BULK twin set (BULK ⊂ FAST). Dash entries denote counts defined relative to the complementary mode; Jaccard overlap reported between the two mode-specific twin sets.

In this benchmarking, all BULK pairs were recovered by FAST mode, while FAST additionally returned 2,778 borderline pairs. This reflects the fact that FAST occasionally selects a slightly smaller *γ* due to the random sub-sample, broadening the kernel and admitting a small number of borderline pairs that fall just below threshold under BULK. From a downstream pipeline perspective, the choice between modes is therefore a trade-off between exact reproducibility (BULK) and broader detection (FAST), not between accuracy and noise. The negative control behavior of FAST closely mirrors that of BULK (median = 0 across three permutations in both modes; BULK worst case: 165 pairs, 0.94% of 17,503; FAST worst case: 200 pairs, 0.99% of 20,281), suggesting that the additional FAST detections are not dominated by obvious permutation-derived artefacts. The worst-case spurious-pair rates across both operating modes remain below 1% of their respective detected-pair totals, collectively serving as an empirical upper bound on the false-positive detection rate and providing post-hoc support for the Z > 2.576 threshold as a practically stringent filter, even in the absence of a formal multiple-testing correction. Recommended use cases for each mode are summarized in Table S2.

### 3.5. Chemical interpretability of detected twins

To illustrate the chemical content of the detected twin pairs, representative examples are shown in Figure 4. The first panel shows a 61-heavy-atom target molecule (a high affinity known BCL2 inhibitor) containing chlorine and sulfur atoms together with its stoichiometric twins; all detected twins maintain similarity scores above 0.985 and preserve the chlorine and sulfur functionality while exhibiting variation in carbon, hydrogen, and oxygen content. The second panel presents a bromine-containing target with multiple perfect-similarity matches (score = 1.000) that alternate between C_14_H_7_BrO_4_ and C_14_H_11_BrO_4_ compositions, illustrating a deliberate property of the current representation. Because hydrogen atom is omitted, molecules that differ only in hydrogen count can receive perfect heavy-atom stoichiometric similarity when their non-hydrogen elemental composition is identical. These examples support that the algorithm retrieves chemically interpretable pairs whose elemental skeletons are conserved while their connectivity is free to vary, which is precisely the property that makes stoichiometric similarity a useful complement to topological fingerprints for scaffold-hopping retrieval.

**Figure 4.**
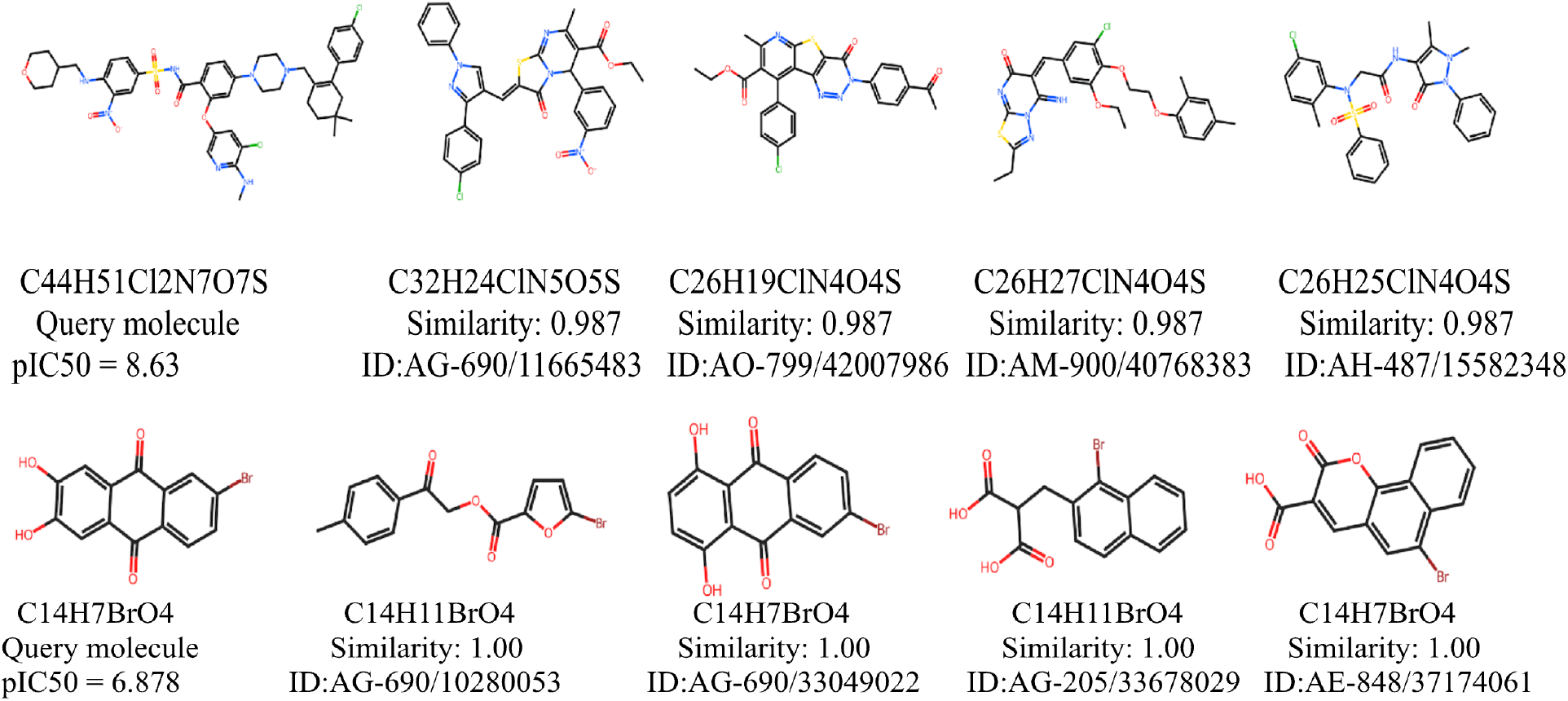
Representative examples of TwinSAR-detected stoichiometric twins, illustrating the chemical interpretability of the algorithm. (Top panel) A 61-heavy-atom target (ID 145, C_44_H_51_Cl_2_N_7_O_7_S, query molecule) and its top-4 stoichiometric twins; all detected twins share similarity scores > 0.985 and preserve the characteristic chlorine and sulfur atoms while permitting variation in carbon, hydrogen, and oxygen content. (Bottom panel) A bromine-containing target (ID 290, C_14_H_7_BrO_4_, query molecule) and its 4 stoichiometric twins; multiple twins achieve perfect similarity scores (1.000) with alternating C_14_H_7_BrO_4_ and C_14_H_11_BrO_4_ compositions, demonstrating that the current heavy-atom representation treats molecules with identical non-hydrogen elemental composition as perfect stoichiometric twins, even when their hydrogen counts differ. The algorithm retrieves pairs whose elemental skeletons are conserved while connectivity is free to vary, the property that makes stoichiometric similarity a useful complement to topological fingerprints for scaffold-hopping retrieval.

### 3.6. BCL-2 case study

As a demonstration of the algorithm’s utility in a realistic virtual screening setting, TwinSAR was deployed in an end-to-end pipeline against the BCL-2 protein, a high-value oncology target whose validated inhibitor Venetoclax (ABT-199) provides a clinically relevant chemical reference.^24,25^ The 692 high-affinity (pIC_50_ >6.50) ChEMBL-BCL2 query molecules were searched against the 327,071-compound SPECS library; TwinSAR retrieved 17,503 twin pairs spanning 10,387 unique SPECS molecules. The relaxed drug-likeness filter retained 99.96% of these (10,383 / 10,387), an empirical confirmation that stoichiometric retrieval inherently operates within biologically relevant chemical space when the queries are themselves drug-like. CatBoost-based pIC_50_ prediction (test *R*^2^ = 0.848; Figure S4 and Table S4) at a pIC_50_ > 6.5 cutoff reduced the candidate pool to 390 molecules, a cumulative 839× cumulative reduction from the original library.

Boltz-2 predicted sequence-dependent changes in BCL-2 pocket accessibility: the truncated model placed the helix over the BH3-binding groove, partially closing the pocket, whereas the full-length model positioned the helix away from the binding site, leaving the groove accessible for ligand binding. Boltz-2 structure-based binding validation against full-length and truncated BCL-2 conformers identified 125 (32.1%) and 105 (26.9%) successful binding hits, respectively (top-10 lists in Figure S6.1 and Figure S6.2, full pipeline statistics in Table S4).

Figure 5 illustrates the chemical content of the top-ranked candidate, C_30_H_29_ClN_4_O_4_S, in relation to its stoichiometric twins from ChEMBL-BCL2. The candidate is structurally distinct from Venetoclax, it presents a triazine-fused benzodiazepine core absent from Venetoclax, yet is stoichiometrically twinned with eleven ChEMBL inhibitors that all carry experimentally measured pIC_50_ values (mean experimental pIC_50_ = 9.81; full twin statistics in Tables S5–S8). Several of these twins (notably ChEMBL Query 631, pIC_50_ = 10.43) display the diagnostic structural motifs of Venetoclax: a chlorinated aromatic ring system, a piperazine/piperidine linker connected to a sulfonamide moiety, a nitro-substituted aromatic ring, and a bulky hydrophobic core comprising multiple fused rings. This outcome, a stoichiometrically related but topologically distinct candidate, retrieved alongside near-neighbors that recapitulate the pharmacophore of a clinically validated drug, is exactly the kind of scaffold-hopping retrieval that motivated the TwinSAR design and provides post-hoc support of the chemical interpretability of the algorithm. Full BCL-2 case study results, including the cumulative reduction cascade, top-10 hit tables, and per-candidate twin profiles, are reported in Section S3 (Supporting Information).

**Figure 5.**
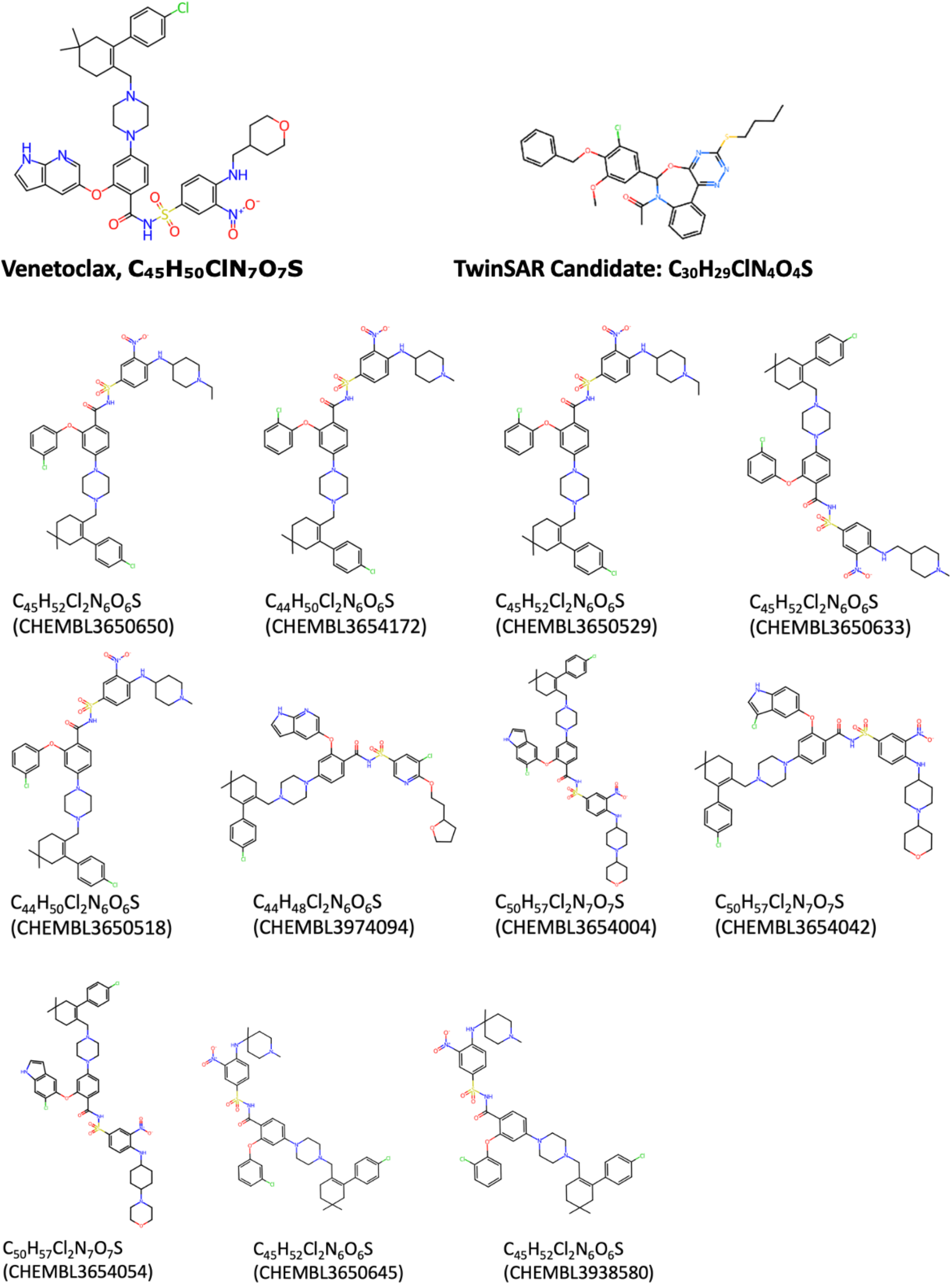
Demonstration of utility (BCL-2 case study). Structural comparison of the top-ranked TwinSAR candidate Specs ID: AN-655/14766010, (C_30_H_29_ClN_4_O_4_S) (top right) with Venetoclax (ABT-199; C_45_H_50_ClN_7_O_7_S, top left) and the eleven stoichiometric twins of the AN-655/14766010 from the ChEMBL-BCL2 dataset. All twins share similarity scores in the range 0.985–0.989 and have a mean experimental pIC_50_ of 9.81. The candidate is topologically distinct from Venetoclax (presenting a triazine-fused benzodiazepine core) but its stoichiometric twins recapitulate the diagnostic Venetoclax pharmacophore: a chlorinated aromatic ring, a piperazine/piperidine linker bearing a sulfonamide, a nitro-substituted aromatic ring, and a bulky multi-ring hydrophobic core.

The top-ranked compound retrieved from the SPECS library by TwinSAR, AN-655/14766010, was modeled in complex with BCL-2 using Boltz-2. (Figure 6) The predicted binding pose indicates that the ligand establishes interactions with known critical amino acid residues within the BCL-2 binding pocket, suggesting that the compound occupies a pharmacologically relevant region of the receptor and may recapitulate key interaction patterns observed for established BCL-2 binders.

**Figure 6.**
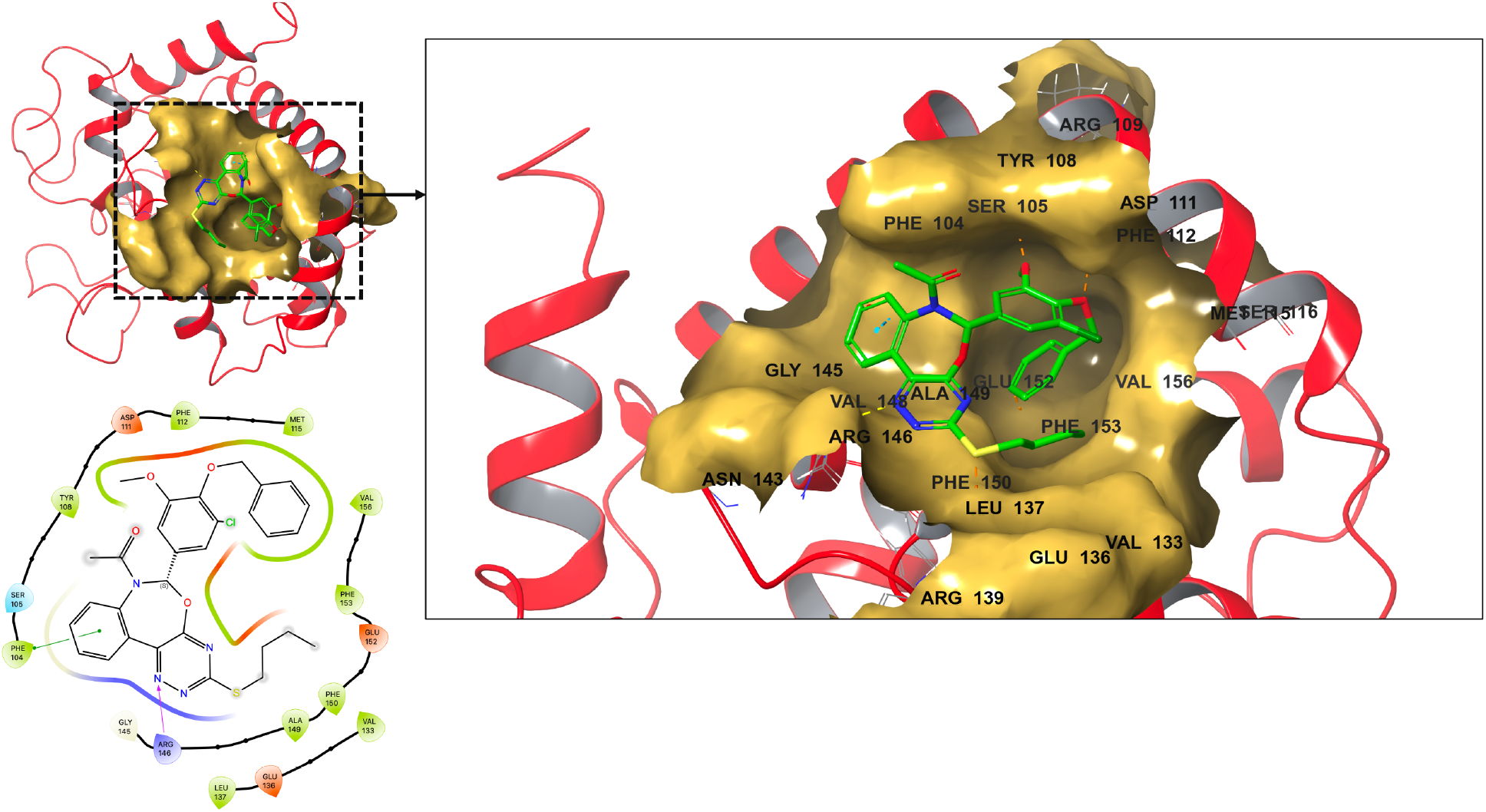
Predicted 3D structure of the BCL-2/AN-655/14766010 complex generated by Boltz-2, with the corresponding 2D ligand interaction map shown alongside.

## 4. Discussion

The present study introduces TwinSAR as a stoichiometry-centered, topology-orthogonal retrieval strategy for large-scale chemical library prioritization. The central observation emerging from the validation experiments is that heavy-atom elemental ratio space contains sufficient information to identify chemically coherent neighborhoods that are not explicitly defined by molecular connectivity. Detected twin pairs were substantially closer in absolute ratio space than block-matched random pairs, support that TwinSAR does not simply retrieve molecules sharing the similar element-presence fingerprint, but instead selects the extreme proximity region within each elemental-composition block. This result supports the use of stoichiometric proximity as a meaningful, computationally inexpensive descriptor layer for early-stage virtual screening.

A key methodological feature of TwinSAR is the combination of exact element-presence blocking with block-wise adaptive RBF scoring. The blocking step provides an efficient way to avoid chemically uninformative comparisons between molecules with incompatible elemental compositions, while reducing the effective all-pairs comparison burden. The adaptive RBF kernel then addresses the fact that different elemental blocks occupy ratio space at different densities. A global kernel width would either over-compress dense blocks or under-resolve sparse ones. By calibrating γ independently within each block, TwinSAR assigns similarity relative to the local stoichiometric environment rather than relative to a single global distance scale. Rather than applying a single global similarity cutoff, the logit-Z filter evaluates each pair relative to the score distribution of its own fingerprint block. This allows chemically dense and sparse blocks to be treated on a comparable statistical scale. This is important because a high raw RBF similarity does not necessarily have the same statistical meaning in all blocks. In a dense block, many molecules may be close to one another, whereas in a sparse block a similar raw score may indicate a much more exceptional match. The moderate relationship between raw similarity and Z-score therefore reflects the intended effect of block normalization. Z-score is not an independent chemical descriptor, but a measure of how extreme a given similarity is relative to the distribution of comparisons within the corresponding block. At the same time, the Gaussian interpretation of the transformed scores should be regarded as approximate. It must be also noted that, although the Z > 2.576 threshold provides a stringent empirical filter, we do not interpret it as a formal false-discovery-rate control because the candidate pairs are not independent and no multiple-testing correction is applied. Instead, the column-permutation negative control provides a dataset-specific empirical estimate of the false-positive rate (worst-case: 0.94% of detected pairs), which we regard as a more contextually appropriate characterization of filter stringency than a parametric correction that would require independence assumptions the data do not satisfy. Empirical per-block calibration of the Z > 2.576 threshold (Supporting Information) supports that it operates conservatively across the majority of fingerprint blocks, though the normal approximation underlying the nominal 0.5% rate is imperfect, as demonstrated by formal normality testing. Thus, this further supports interpreting the threshold as a practically calibrated empirical filter rather than a parametric statistical guarantee.

The representation comparison provides an important simplification. The 8-element single-ratio representation and the 254-feature subset-ratio representation produced identical twin-pair sets in the controlled benchmark, while the higher-dimensional representation incurred a substantial runtime penalty. This equivalence is chemically and mathematically interpretable: in conventional drug-like organic molecules, the relevant stoichiometric information is already captured by the single-element heavy-atom ratios, whereas subset-ratio features are deterministic functions of the same elemental counts. Thus, the *single8* representation offers the best trade-off between sensitivity, speed and interpretability for the chemical space examined here. The *full254* representation may still be useful in specialized chemical domains. The comparison between BULK and FAST modes demonstrates the practical flexibility of the algorithm. In the present production benchmark, FAST mode reduced wall-clock time while retaining the same unique-query and unique-target coverage as BULK and adding additional borderline pair-level matches. This suggests that stochastic γ estimation may be suitable for exploratory, recall-prioritized screening campaigns. However, because FAST relies on random sampling of intra-block distances, the observed nesting of BULK pairs within FAST pairs should be interpreted as an empirical property of the present dataset rather than a formal guarantee.

The BCL-2 case study illustrates how TwinSAR can be embedded into a multi-stage virtual screening workflow. Starting from a large commercial library (>300.000 compounds), the algorithm reduced the search space to a focused set of stoichiometrically related candidates, which were subsequently filtered by drug-likeness, prioritized using a CatBoost potency model and evaluated by structure-based binding probability prediction. This workflow demonstrates the practical utility of TwinSAR as a candidate-set compression tool.

Several limitations of the present algorithm should also be noted. The TwinSAR retrieves molecules that conserve heavy-atom stoichiometric skeletons while allowing connectivity to vary. This property is valuable because it provides a route to topology-orthogonal candidate expansion. Because hydrogen is deliberately omitted, compounds differing only in hydrogen count can be assigned perfect stoichiometric similarity when their heavy-atom composition is identical. This is a design choice that improves robustness to protonation and tautomer assignment, but it also reduces sensitivity to saturation state and valence-related differences. The current eight-element implementation is optimized for conventional medicinal chemistry and may require extension for phosphorus-, boron-, silicon-, selenium- or metal-containing libraries.

Despite these limitations, TwinSAR occupies a useful methodological niche. It is fast, interpretable, conformation-free and orthogonal to widely used fingerprint metrics. Its greatest utility is likely to be as an early-stage prefilter that broadens candidate discovery beyond close structural analogues while still preserving a chemically meaningful composition envelope. In the future work, we will focus on soft element-class blocking, weighted or atom-type-aware ratio representations, empirical false discovery rate (FDR) calibration, GPU acceleration and systematic retrospective benchmarking across multiple target classes. Integration with ECFP/Tanimoto, pharmacophore, shape and docking workflows may also provide a more complete strategy for scaffold-diverse hit discovery than any individual similarity metric alone.

## 5. Conclusions

We have introduced TwinSAR, a stoichiometric twin-detection algorithm whose novelty resides in three coordinated methodological innovations. First, binary fingerprint blocking partitions molecules by element-presence patterns and reduces the cost of all-pairs comparison from O(*NM*) to *O*(∑*n*_*i*_*m*_*i*_), enabling stoichiometric searches at million-billion compound scale. Second, a per-block adaptive RBF kernel calibrated by the median heuristic provides fair similarity comparison across chemical sub-spaces of vastly different density; the substantial block-to-block variation of γ across blocks demonstrates the necessity of this adaptation. Third, a logit-transformed Z-score filter maps bounded RBF scores onto an unbounded scale and provides a stringent block-wise filtering layer for prioritizing statistically exceptional stoichiometric pairs.

Statistical validation across three orthogonal tests supports the algorithmic soundness of TwinSAR: Detected twin pairs are 12.7× more similar than block-matched random pairs (*p* < 0.001, a column-permutation negative control returns a median of zero spurious twins), and in the present benchmark, FAST recovered all BULK pairs while providing a 3.29× speed-up, although this nesting should be interpreted as an empirical observation. A controlled benchmark further established that the streamlined 8-element representation is sensitivity-equivalent to a comprehensive 254-element representation (Jaccard = 1.000 on detected twins) while running 3.55× faster, supporting its adoption as the production default. Inspection of representative detected twin pairs showed that the algorithm retrieves chemically interpretable structures whose elemental skeletons are conserved while connectivity is free to vary, precisely the property that distinguishes scaffold-hopping retrieval from connectivity-based approaches.

TwinSAR was also embedded in an end-to-end virtual screening pipeline against the BCL-2 target protein and reduced a 327,071-compound commercial library by 839-fold to a focused candidate panel; the top-ranked candidate is stoichiometrically twinned with sub-nanomolar ChEMBL binders that recapitulate the Venetoclax pharmacophore. TwinSAR offers a fast, conformation-free, and statistically principled prefilter for stoichiometric similarity that is fully orthogonal to topological fingerprints. Because the algorithm is target- and pipeline-agnostic, integration of TwinSAR with established Tanimoto-based searches and pharmacophore screens should materially improve the recall of scaffold-hopping bioisosteres in modern hit-discovery campaigns. Future methodological work will focus on (i) extending the algorithm to weighted ratio representations that incorporate atom-typing information; (ii) exploring kernel choices beyond the Gaussian RBF; (iii) GPU acceleration for the kernel-evaluation step, which dominates the compute budget at large block sizes; and (iv) systematic comparison with topological-fingerprint and shape-matching baselines on standard scaffold-hopping benchmarks.

Beyond its performance in the BCL-2 screening workflow, the principal innovation of TwinSAR lies in redefining molecular similarity as a statistically normalized stoichiometric relationship rather than a purely topology-dependent property. By integrating exact element-presence blocking, per-block adaptive RBF kernel calibration, and logit-transformed Z-score filtering into a single retrieval framework, TwinSAR addresses three major limitations of conventional similarity searching: inefficient all-pairs comparison, non-uniform chemical-space density, and the poor statistical comparability of raw bounded similarity scores. This design allows each candidate pair to be evaluated not only by its absolute elemental-ratio proximity, but also by how exceptional that proximity is within its own elemental-composition background. The finding that the compact 8-element heavy-atom representation reproduced the 254-feature representation without loss of sensitivity further demonstrates that chemically meaningful stoichiometric information can be captured in a minimal, interpretable, and computationally efficient descriptor space. Importantly, TwinSAR introduces an orthogonal prefiltering layer capable of enriching scaffold-diverse candidates that may be missed by connectivity-based metrics. This combination of scalability, statistical normalization, chemical interpretability, and topology independence represents the most distinctive methodological contribution of the present study and supports the use of stoichiometric twin detection as a complementary strategy in large-scale virtual screening and scaffold-hopping-oriented hit discovery.

## Supporting Information

The Supporting Information document accompanying this manuscript contains: Section S1, BCL-2 protein FASTA sequences used for Boltz-2 binding validation; Section S2, CatBoost optimal hyperparameters and reproducibility artifacts; Section S3, detailed BCL-2 case study results; Figure S1, the effect of logit transformation on score distribution; Figure S2, a five-panel diagnostic suite for normality and block-size reliability; Figure S3, algorithmic validation of TwinSAR in FAST mode; Figure S4, CatBoost regression model diagnostics; Figure S5, permutation feature importance for molecular descriptors; Figure S6.1, top-10 hits ranked by pIC_50_ with vs. without the C-terminal helix; Figure S6.2, a side-by-side comparison of predicted pIC_50_ values; Table S1, normality test statistics for transformed Z-scores; Table S2, column-permutation negative-control results; Table S3, recommended TwinSAR processing modes; Table S4, the multi-tier BCL-2 library compression cascade; and Tables S5–S8, molecule–twin summary tables for the top-ranked BCL-2 candidates.

## Conflicts of interest

The authors declare no competing financial interest.

## Data and code availability

The TwinSAR source code and scripts are available at https://github.com/DurdagiLab/Twinsar. The ChEMBL-BCL2 dataset is freely available from www.ebi.ac.uk/chembl. The SPECS commercial library is available from www.specs.net.

## Notes

### Competing Interest Statement

The authors have declared no competing interest.

