## Supporting Material for "TwinSAR: An Adaptive Kernel-based Algorithm with logit-transformed Z-score Filtering for Chemical Twin Detection in Large-scale Virtual Screening"

TwinSAR
Kernel-Based Molecular Twin Discovery

### S1. BCL-2 Protein FASTA Sequences for Boltz-2 Binding Validation

Two BCL-2 receptor inputs were used in the Boltz-2 binding validation step of the BCL-2 case study. The full-length sequence (residues 1–239) includes the C-terminal helix that anchors BCL-2 to the outer mitochondrial membrane. The truncated sequence (residues 1–206) omits this helix and corresponds to common crystallographic constructs.

#### Full length (with C-terminal helix):

```
MAHAGRTGYDNREIVMKYIHYKLSQRGYEWDAGDVGAAPPGAAPAPGIFSS-
QPGHTPHPAASRDPVARTSPLQTPAAPGAAAGPALSPVPPVVHLTLRQAGDDFSRRYRRDFAEMSSQLHLTPFTARGRFATVVEEL
FRDGVNWGRIVAFFEFGGVMCVESVNREMSPLVDNIALWMTEYLNRHL-
HTWIQDNGGWDAFVELYGPSMRPLFDFSWLSLKTLLSLALVGACITLGAYLGHK
```

#### Truncated (without C-terminal helix):

```
MAHAGRTGYDNREIVMKYIHYKLSQRGYEWDAGDVGAAPPGAAPAPGIFSS-
QPGHTPHPAASRDPVARTSPLQTPAAPGAAAGPALSPVPPVVHLTLRQAGDDFSRRYRRDFAEMSSQLHLTPFTARGRFATVVEEL
FRDGVNWGRIVAFFEFGGVMCVESVNREMSPLVDNIALWMTEYLNRHLHTWIQDNGGWDAFVELYGPSMR
```

### S2. CatBoost Optimal Hyperparameters and Reproducibility Artefacts

All artefacts required for reproduction of the CatBoost $pIC_{50}$ predictor used in the BCL-2 case study. The optimal hyperparameter set identified by the three-fold cross-validated grid search (108 configurations) is summarized below. The complete preprocessing pipeline (variance threshold selector, median imputer, MinMaxScaler) and the final regressor trained on all 1,017 ChEMBL-BCL2 molecules with experimental $pIC_{50}$ are serialized in joblib format. The optimized CatBoost regressor achieved test $R^2$ = 0.848 and RMSE = 0.738 with a negligible validation–testset gap ($\Delta R^2$ = 0.011), indicating no significant overfitting despite the exhaustive grid search.

#### Optimal CatBoost hyperparameters:

| Hyperparameter | Optimal value |
|---|---|
| depth | 8 |
| iterations | 300 |
| learning_rate | 0.03 |
| l2_leaf_reg | 1 |
| loss_function | RMSE |
| random_state | 42 |

### S3. BCL-2 Case Study Results

The complete library compression cascade is shown in Table S4, the Boltz-2 top-10 hit molecules under the two helix conformations are illustrated in Figure S6.1, the side-by-side comparison of predicted $pIC_{50}$ between conformations is shown in Figure S6.2, and detailed molecule–twin summary tables for the top four BCL-2 candidates are provided in Tables S5–S8.

Out of the 390 high-confidence candidates submitted to Boltz-2, 125 (32.1%) crossed the binding-probability threshold $P_{\mathrm{bind}} > 0.5$ against full-length BCL-2 (with C-terminal helix), and 105 (26.9%) qualified as hits against the truncated construct (without helix). The same molecule, $C_{30}H_{29}ClN_4O_4S$, ranked #1 in both conditions, with predicted $pIC_{50}$ of 7.273 (with helix) and 7.192 (without). Inspection of the predicted complexes revealed that the helix folded into a closed conformation against the BH3 groove for approximately half of the cases and remained extended for the rest, consistent with the known dynamic disorder of this transmembrane segment in solution.

### Supporting Figures

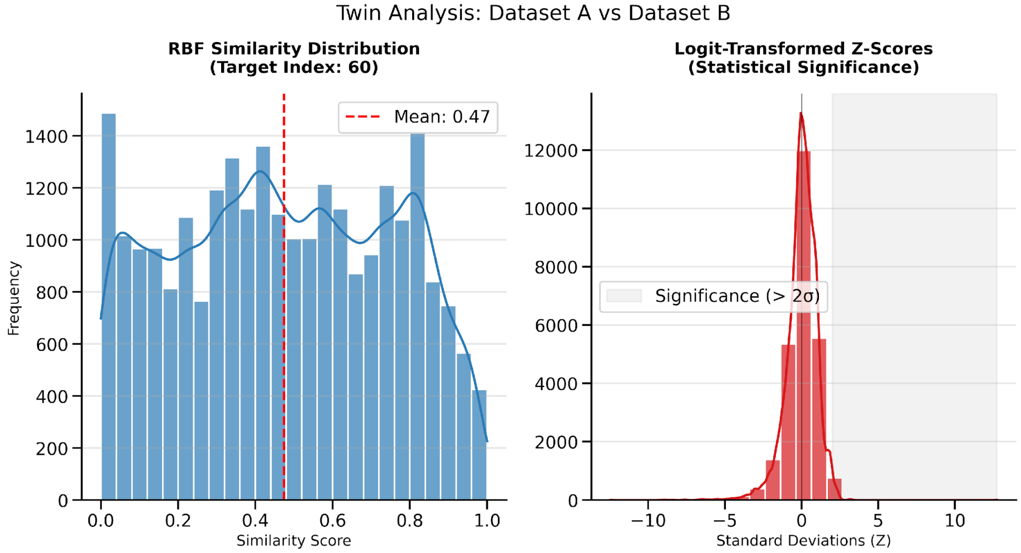


**Figure S1.** Effect of the logit transformation on the score distribution (BULK mode, 8-element representation, single representative target). (Left) Raw RBF similarity distribution for the target's neighbourhood; the strong negative skew compresses true twins near similarity = 1.0 and prevents direct Gaussian thresholding. (Right) Logit-transformed Z-score distribution for the same data; the values are approximately normally distributed about zero, allowing the application of standard Z-score statistics. The $Z > 2.576$ threshold (one-sided $p < 0.005$) used by TwinSAR retains only the extreme right-hand tail of the distribution, as visible in the histogram.

### Normality Diagnostics of Logit-Transformed Z-Scores

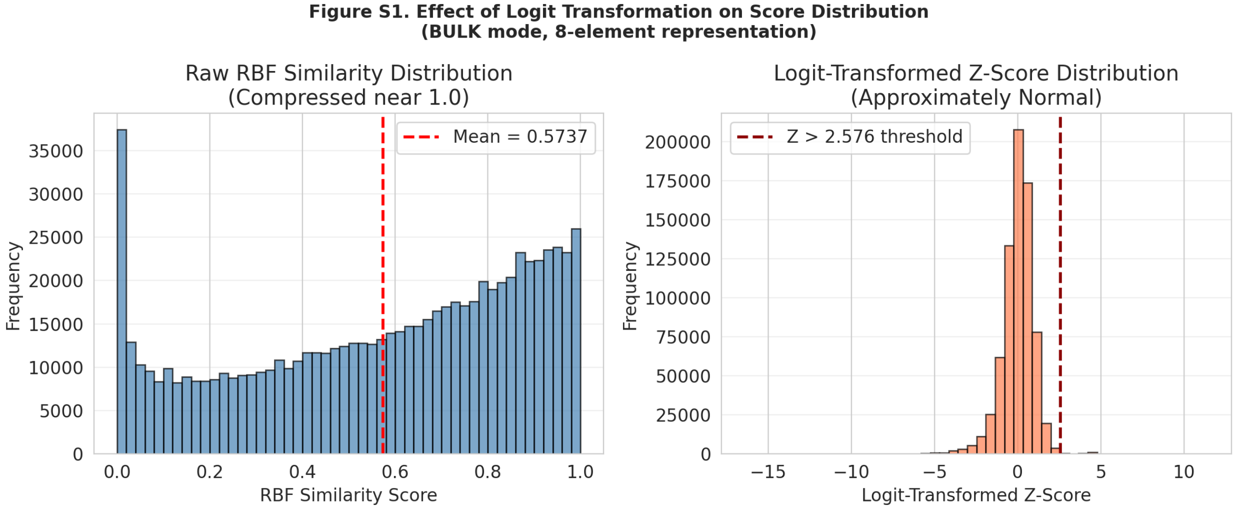


**Figure S2.1.** Effect of the logit transformation on the score distribution (BULK mode, 8-element representation, single representative target). (Left) Raw RBF similarity distribution for the target's neighbourhood; the strong negative skew compresses true twins near similarity = 1.0 and prevents direct Gaussian thresholding. (Right) Logit-transformed Z-score distribution for the same data; the values are approximately normally distributed about zero, allowing the application of standard Z-score statistics. The $Z > 2.576$ threshold (nominally corresponding to the upper 0.5% tail of the standard normal distribution, applied here as an empirical block-wise filter criterion) retains only the extreme right-hand tail of the distribution, as visible in the histogram. Formal normality tests (Kolmogorov–Smirnov, Shapiro–Wilk) rejected the normal null hypothesis for all blocks ($p < 0.001$ in all cases), reflecting the characteristic negative skewness and excess kurtosis of the logit-transformed RBF score distributions (see Normality Test Statistics table and Q–Q plots). Empirical per-block false-positive rates at $Z = 2.576$ were below the nominal 0.5% in the majority of blocks, confirming that the threshold operates conservatively across most of the chemical space examined; block-level departures above the nominal rate were confined to a minority of structurally compact blocks with high kurtosis.

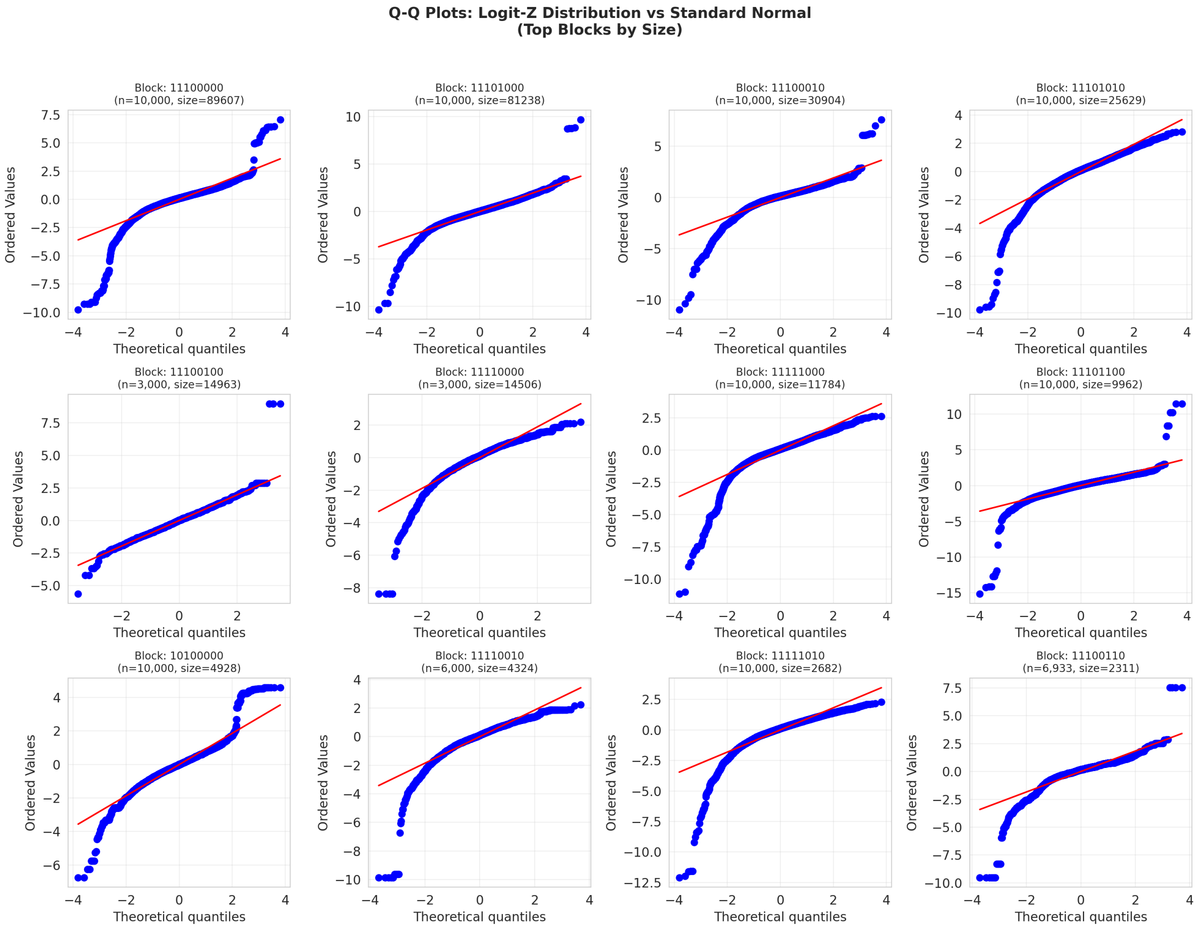


**Figure S2.2.** Q-Q plots of logit-transformed Z-scores versus standard normal distribution for the top12 fingerprint blocks by size. Each panel shows the deviation from normality for a specific block, with block identifier, sample size (n), and block size (number of targets) noted. Departures from the diagonal line indicate non-normality, consistent with the formal test results. The characteristic S-shaped deviations reflect the negative skewness and excess kurtosis noted in the main text.

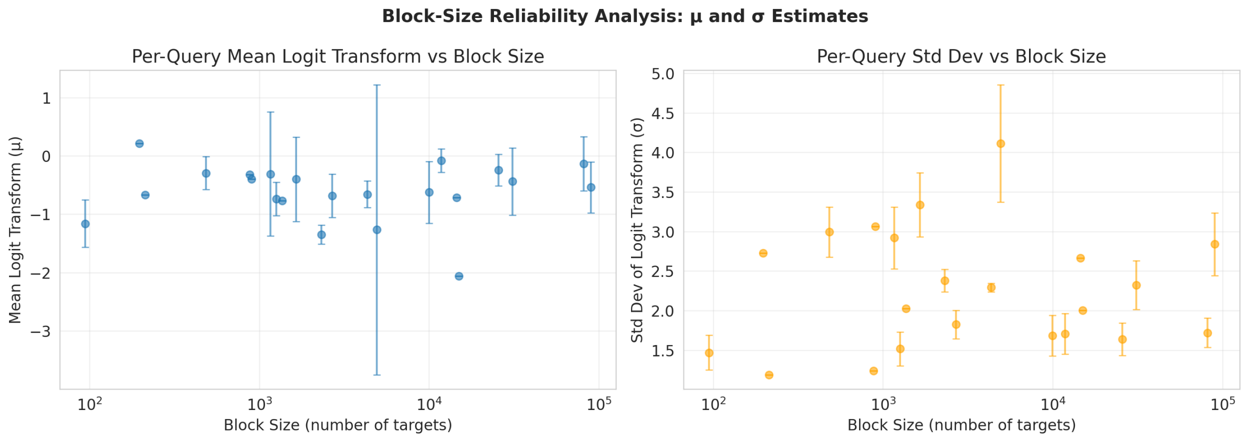


**Figure S2.3.** Block-size reliability analysis of μ and σ estimates. (Left) Mean logit transform (μ) per query versus block size, with error bars indicating inter-query standard deviation. (Right) Standard deviation of logit transform (σ) per query versus block size. Block sizes span two orders of magnitude (~10 to ~1000 targets). The estimates remain stable across the range, confirming that the per-block adaptive kernel is robust to block size variation.

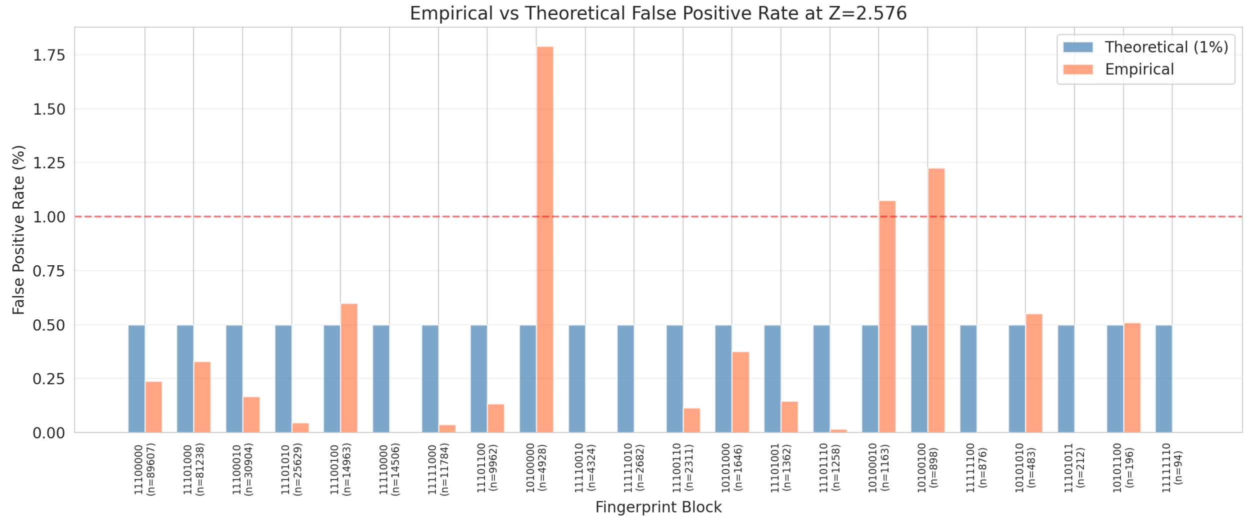


**Figure S2.4.** Empirical versus theoretical false positive rate at Z = 2.576 threshold across fingerprint blocks. Blue bars show the nominal 1% rate (theoretical, based on standard normal approximation). Coral bars show the empirically observed false positive rate per block. Most blocks exhibit empirical rates below the nominal rate, confirming conservative threshold behavior. The minority of blocks exceeding the nominal rate correspond to structurally compact blocks with high kurtosis, consistent with the Q-Q plot deviations and formal normality test results.

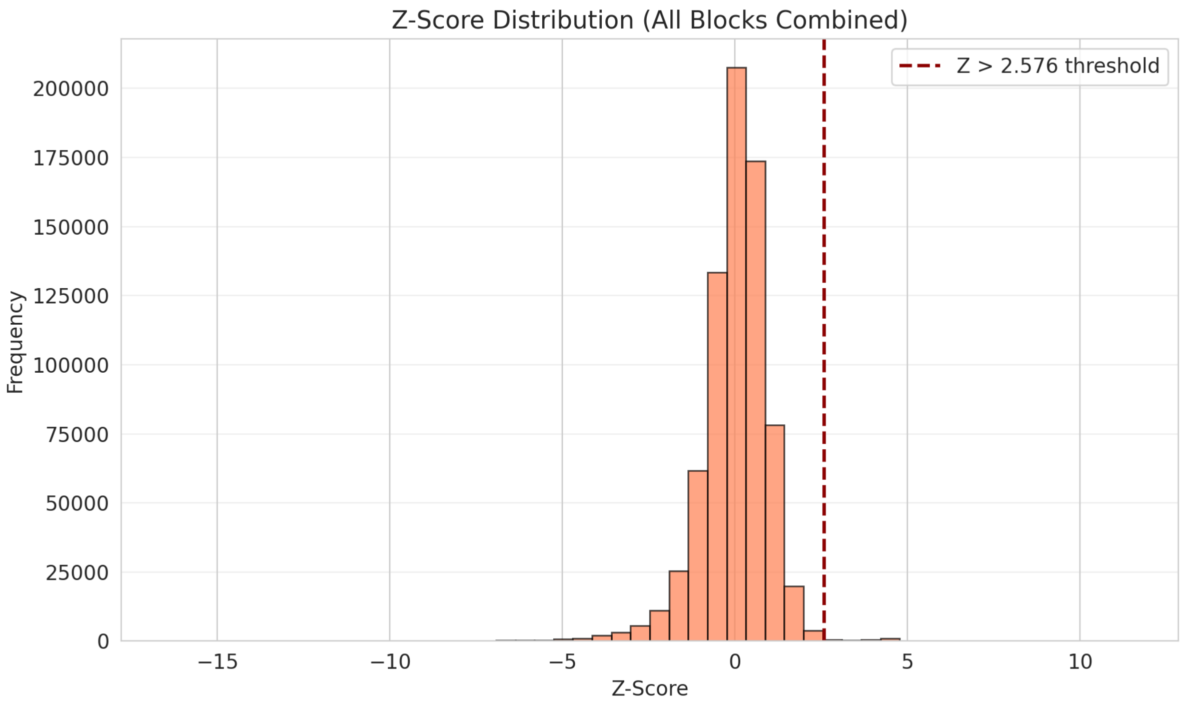


**Figure S2.5.** Z-score distribution across all fingerprint blocks. The histogram shows the combined logit-transformed Z-scores from all blocks, with the Z > 2.576 threshold marked (dashed line). The approximately normal distribution about zero enables block-wise statistical filtering; the heavy right-hand tail corresponds to statistically exceptional stoichiometric twins retained by TwinSAR.

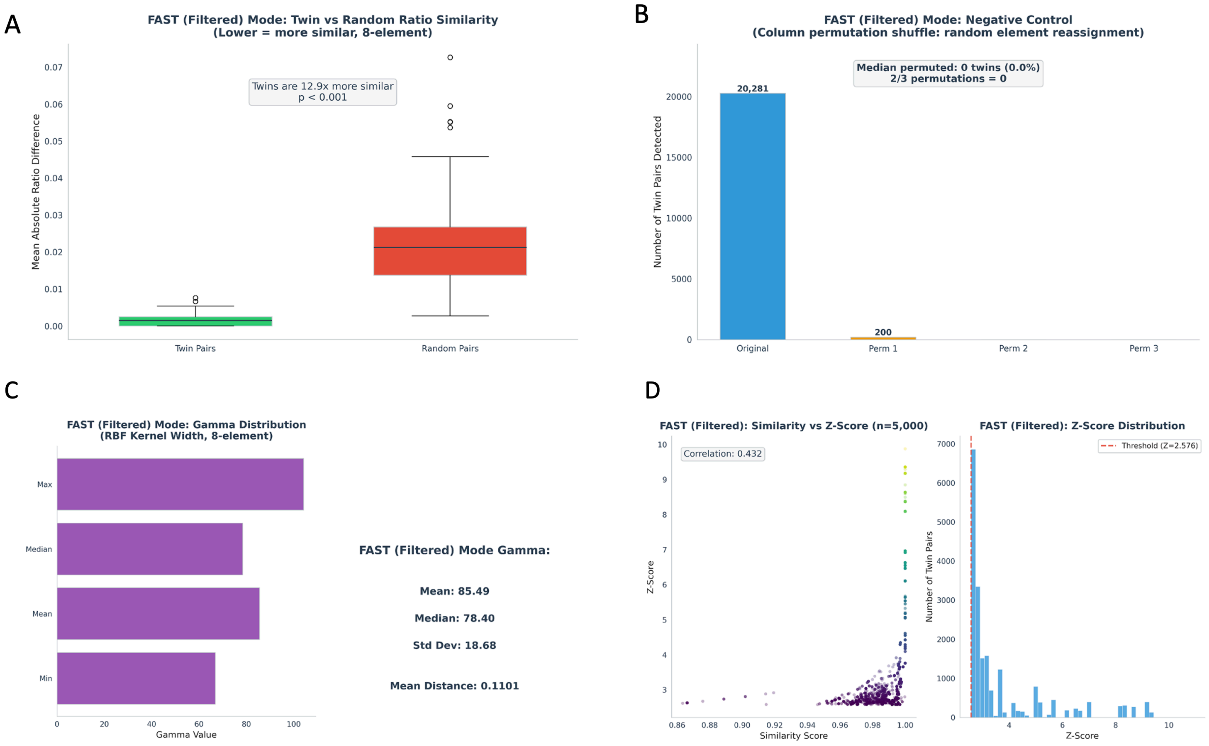


**Figure S3.** Algorithmic validation of TwinSAR in FAST mode (8-element representation, $pIC_{50} > 6.5$ pre-filtered query set). (A) Twin pairs are 12.7× more similar in ratio space than random pairs from the same fingerprint block ($p < 0.001$). (B) Negative control via column-permutation shuffle: median permuted twins = 0, confirming that FAST mode also rejects scrambled element assignments. (C) Distribution of the adaptive kernel precision parameter $\gamma$ across fingerprint blocks (mean = 85.49, median = 78.40), identical to BULK mode. (D) Per-target similarity vs. Z-score relationship (Pearson $r = 0.432$). The FAST-mode validation profile mirrors that of BULK mode (Figure 2, main text), confirming that the 3.29x wall-clock speed-up does not compromise statistical behavior.

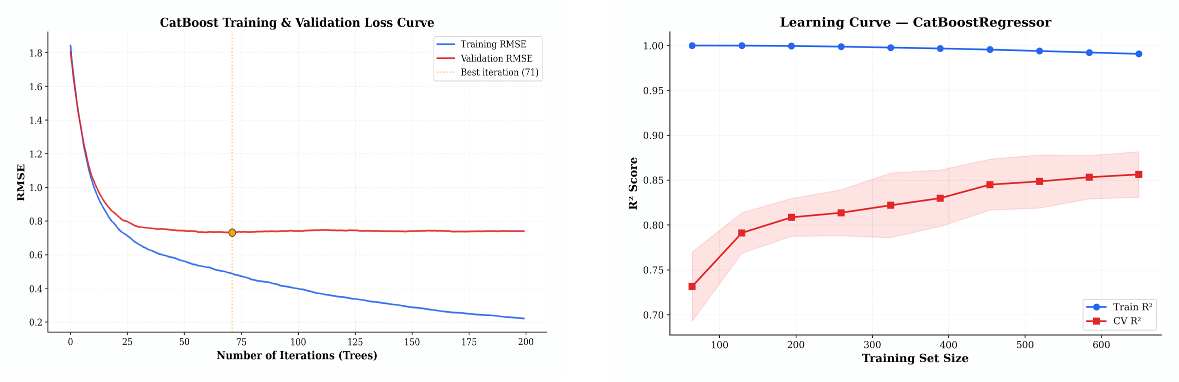


**Figure S4.** CatBoost $pIC_{50}$ regression model diagnostics for BCL-2 case study. (A) Training (blue) and validation (red) RMSE curves over 300 boosting iterations. (B) Learning curves on five-fold cross-validation across progressively larger training subsets demonstrate convergence of training and cross-validation $R^2$, indicating adequate model complexity and low variance.

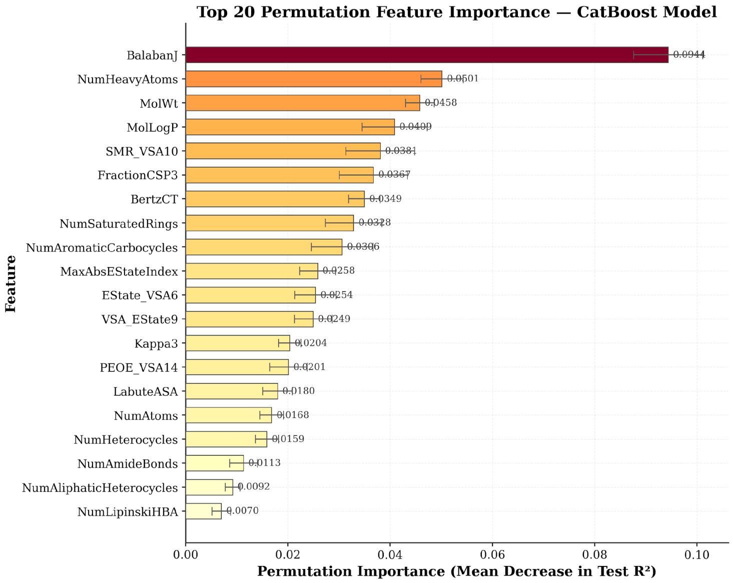


**Figure S5.** Top 20 molecular descriptors ranked by permutation importance (mean decrease in test $R^2$). The topological connectivity index BalabanJ dominates ($\Delta R^2 = 0.098$), followed by NumHeavyAtoms, MolWt, and MolLogP. The model achieved test $R^2 = 0.848$ and RMSE = 0.738.

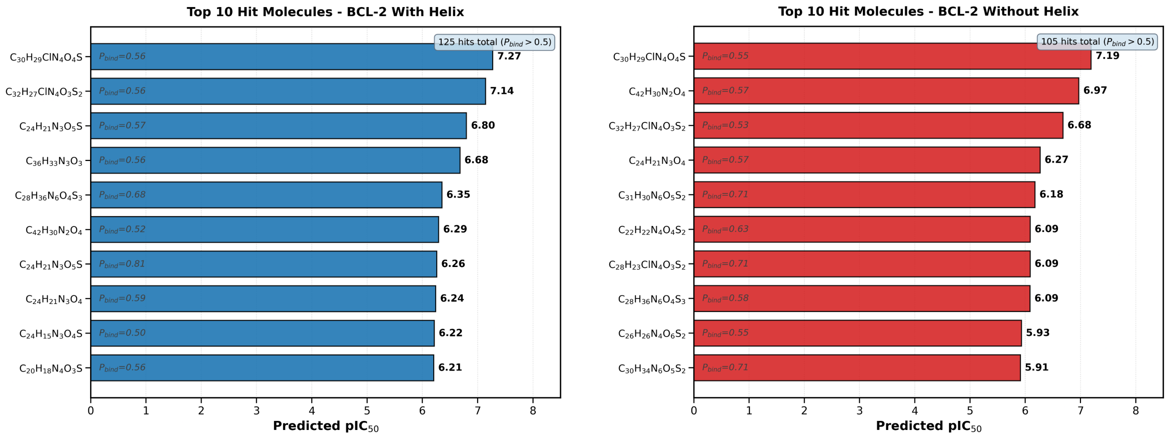


**Figure S6.1.** Top 10 hit molecules ranked by predicted binding affinities from Boltz-2 validation in the BCL-2 case study. (Left) BCL-2 with C-terminal helix retained (blue bars). (Right) BCL-2 with C-terminal helix removed (red bars). All molecules shown satisfy the binding-probability threshold ($P_{bind} > 0.5$). Numerical labels above each bar indicate the Boltz-2 binding probability and the predicted $pIC_{50}$. The molecule $C_{30}H_{29}ClN_4O_4S$ ranks #1 in both conditions, demonstrating helix-independent prioritization; molecules in the lower portion of each list show condition-specific ranking.

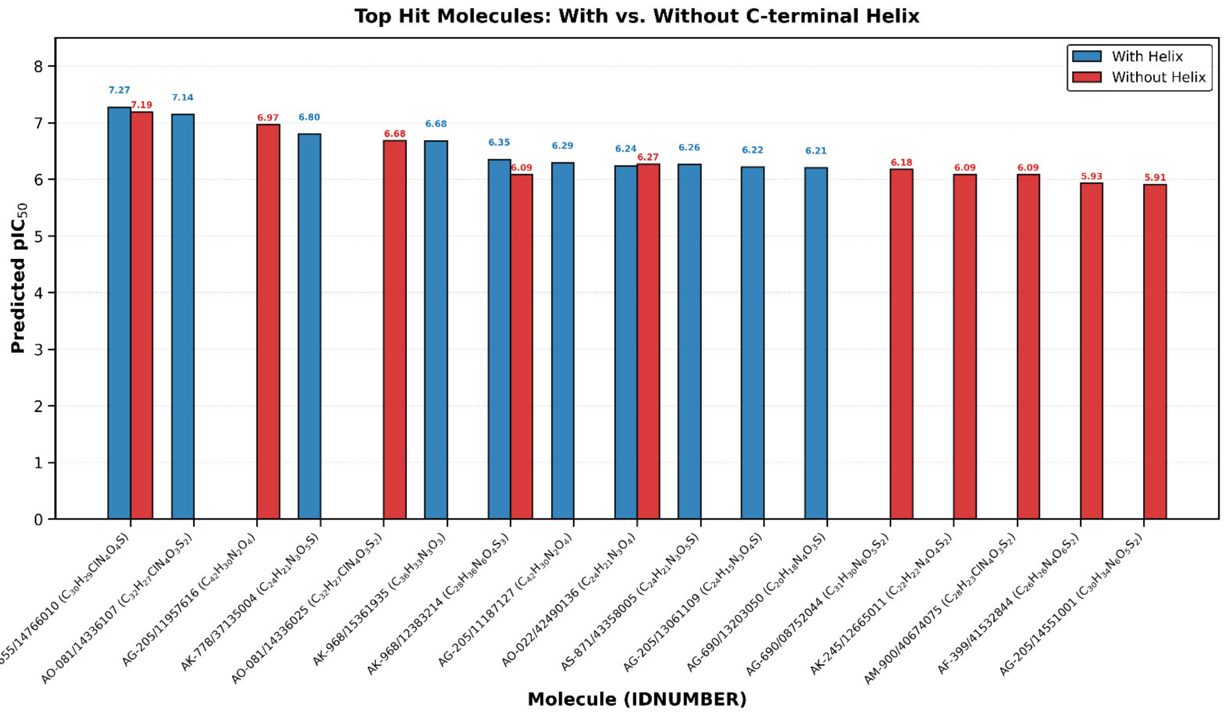


**Figure S6.2.** Side-by-side comparison of Boltz-2 predicted binding affinities for all unique molecules appearing in either top-10 hit list (with helix or without helix), identified by chemical formula. Blue bars = full-length BCL-2 (with C-terminal helix); red bars = truncated BCL-2 (without C-terminal helix).

### Supporting Tables

**Table S1.** Normality test statistics for logit-transformed Z-scores across fingerprint blocks. Skewness and Kurtosis quantify distributional asymmetry and tail weight. Shapiro-Wilk p-values (when available) and Kolmogorov-Smirnov p-values indicate formal rejection of the normal null hypothesis ($p < 0.001$ in all cases). Empirical false positive rate (Emp FP%) is the observed fraction of scores exceeding $Z = 2.576$, compared to the theoretical rate of 0.5%. The threshold operates conservatively (empirical < theoretical) in most blocks, with departures confined to smaller blocks with high kurtosis.

| Block | Size | N | Skewness | Kurtosis | Shapiro p | KS p | Emp FP% | Theo FP% |
|---|---|---|---|---|---|---|---|---|
| 10100000 | 4928 | 90000 | 0.580 | 5.609 | 0.0000 | 0.0000 | 1.79 | 0.50 |
| 10100010 | 1163 | 4652 | 0.101 | 8.105 | 0.0000 | 0.0000 | 1.07 | 0.50 |
| 10100100 | 898 | 898 | 1.020 | 11.765 | 0.0000 | 0.0000 | 1.22 | 0.50 |
| 10101000 | 1646 | 8230 | -1.624 | 11.752 | 0.0000 | 0.0000 | 0.38 | 0.50 |
| 10101010 | 483 | 1449 | -1.442 | 14.391 | 0.0000 | 0.0000 | 0.55 | 0.50 |
| 10101100 | 196 | 196 | 0.274 | 5.557 | 0.0000 | 0.0796 | 0.51 | 0.50 |
| 11100000 | 89607 | 90000 | -1.752 | 14.903 | 0.0000 | 0.0000 | 0.24 | 0.50 |
| 11100010 | 30904 | 90000 | -1.709 | 10.509 | 0.0000 | 0.0000 | 0.17 | 0.50 |
| 11100100 | 14963 | 3000 | 0.595 | 6.570 | 0.0000 | 0.0080 | 0.60 | 0.50 |
| 11100110 | 2311 | 6933 | -1.969 | 14.435 | 0.0000 | 0.0000 | 0.12 | 0.50 |
| 11101000 | 81238 | 90000 | -0.778 | 7.071 | 0.0000 | 0.0000 | 0.33 | 0.50 |
| 11101001 | 1362 | 1362 | -1.685 | 16.076 | 0.0000 | 0.0000 | 0.15 | 0.50 |
| 11101010 | 25629 | 90000 | -1.480 | 7.574 | 0.0000 | 0.0000 | 0.05 | 0.50 |
| 11101011 | 212 | 212 | -0.777 | 1.215 | 0.0001 | 0.0603 | 0.00 | 0.50 |
| 11101100 | 9962 | 69000 | -1.912 | 26.809 | 0.0000 | 0.0000 | 0.13 | 0.50 |
| 11101110 | 1258 | 37740 | -1.034 | 2.253 | 0.0000 | 0.0000 | 0.02 | 0.50 |
| 11110000 | 14506 | 3000 | -1.900 | 8.985 | 0.0000 | 0.0000 | 0.00 | 0.50 |
| 11110010 | 4324 | 6000 | -2.448 | 16.576 | 0.0000 | 0.0000 | 0.00 | 0.50 |
| 11111000 | 11784 | 57000 | -2.113 | 11.483 | 0.0000 | 0.0000 | 0.04 | 0.50 |
| 11111010 | 2682 | 80460 | -2.668 | 18.892 | 0.0000 | 0.0000 | 0.00 | 0.50 |
| 11111100 | 876 | 876 | -0.987 | 3.833 | 0.0000 | 0.0224 | 0.00 | 0.50 |
| 11111110 | 94 | 564 | -0.617 | 0.455 | 0.0000 | 0.0400 | 0.00 | 0.50 |

**Table S2.** Negative-control results from column-permutation shuffle of the eight ratio columns. BULK mode, 8-element ratio representation, query set pre-filtered to retain ChEMBL-BCL2 molecules with experimental $pIC_{50} > 6.5$ ($n = 692$). Each permutation applies a single global shuffle of the eight element-ratio columns to every target molecule, breaking the chemical identity of each ratio while preserving the per-column value distributions. "Common blocks" denotes the number of fingerprint blocks shared between the permuted target set and the query set. "Fraction" is the twin-pair count expressed as a percentage of the unshuffled baseline (17,503 pairs). FAST mode produced equivalent behaviour (median = 0; 200 pairs from permutation 1, 0.98% of 20,281).

| Dataset | Common blocks | Twin pairs | Fraction | Status |
|---|---|---|---|---|
| Unshuffled (baseline) | 17 | 17,503 | 100 | Reference |
| Permutation 1 | 1 | 165 | 0.0094 | Pass |
| Permutation 2 | 10 | 0 | 0.0 | Pass |
| Permutation 3 | 13 | 0 | 0.0 | Pass |
| Median over permutations | — | 0 | 0.0 | Pass |

**Table S3.** Recommended TwinSAR processing mode by use case. FAST is preferred when wall-clock saving outweighs minor run-to-run variability; BULK is preferred whenever exact reproducibility is required.

| Use case | Recommended mode |
|---|---|
| Exploratory analysis, large-scale screening | FAST |
| Final results for publication | BULK |
| Regulatory submissions | BULK |
| Method development and parameter tuning | FAST |
| Cross-study comparisons / meta-analysis | BULK |
| Maximal candidate detection (recall-priority) | FAST |

**Table S4.** BCL-2 multi-tier library compression cascade (case study). The combination of stoichiometric retrieval (3.2% retention), druglikeness filtering (99.96% retention), and CatBoost potency prediction (3.8% retention of druglike twins) reduced 327,071 SPECS molecules to 390 high-confidence candidates, a cumulative 839× library compression. Boltz-2 then partitioned these 390 candidates into 125 hits (with helix) and 105 hits (without helix).

| Pipeline stage | Compounds | Stage retention | Cumulative library compression |
|---|---|---|---|
| SPECS library | 327,071 | 100% | 1× |
| TwinSAR retrieval (8-elem.) | 10,387 | 3.2% | 31.5× |
| Druglikeness filter | 10,383 | 99.96% | 31.5× |
| CatBoost $pIC_{50} > 6.5$ | 390 | 3.8% | 839× |
| Boltz-2 hits, with helix | 125 | 32.1% | 2,617× |
| Boltz-2 hits, without helix | 105 | 26.9% | 3,115× |

**Table S5.** Top-ranked candidate $C_{30}H_{29}ClN_4O_4S$ (rank #1 in both helix conditions; main text Figure 5) and its top five stoichiometric twins from the ChEMBL-BCL2 reference set, ranked by experimental $pIC_{50}$. Sim. = RBF similarity score; Z = logit-transformed Z-score; Exp. $pIC_{50}$ = experimental $pIC_{50}$ of the twin from ChEMBL.

**Query molecule properties:**

| Property | Value |
|---|---|
| Heavy atoms | 40 |
| Predicted $pIC_{50}$ (with helix / without helix) | 7.273 / 7.192 |
| Boltz-2 $P_{bind}$ (with / without helix) | 0.560 / 0.552 |
| Classification | Hit (both conditions) |
| Number of twins | 11 |
| Mean twin exp. $pIC_{50}$ | 9.811 |
| Median twin exp. $pIC_{50}$ | 10.046 |
| Max twin exp. $pIC_{50}$ | 10.432 |
| Min twin exp. $pIC_{50}$ | 7.548 |

**Top-5 twins by experimental $pIC_{50}$:**

| # | Twin formula | Sim. | Z | Exp. $pIC_{50}$ |
|---|---|---|---|---|
| 1 | $C_{45}H_{52}Cl_2N_6O_6S$ | 0.9885 | 2.670 | 10.432 |
| 2 | $C_{44}H_{50}Cl_2N_6O_6S$ | 0.9862 | 2.597 | 10.310 |
| 3 | $C_{45}H_{52}Cl_2N_6O_6S$ | 0.9885 | 2.670 | 10.301 |
| 4 | $C_{45}H_{52}Cl_2N_6O_6S$ | 0.9885 | 2.670 | 10.208 |
| 5 | $C_{44}H_{50}Cl_2N_6O_6S$ | 0.9862 | 2.597 | 10.155 |

**Table S6.** Candidate $C_{32}H_{27}ClN_4O_3S_2$ (rank #2 with helix, rank #3 without helix) and its top stoichiometric twin.

**Query molecule properties:**

| Property | Value |
|---|---|
| Heavy atoms | 42 |
| Predicted $pIC_{50}$ (with helix / without helix) | 7.142 / 6.684 |
| Boltz-2 $P_{bind}$ (with / without helix) | 0.560 / 0.526 |
| Classification | Hit (both conditions) |
| Number of twins | 1 |

**Top-twin:**

| Twin formula | Sim. | Z | Exp. $pIC_{50}$ |
|---|---|---|---|
| $C_{31}H_{27}ClN_4O_3S_2$ | 0.9964 | 3.103 | 6.891 |

**Table S7.** Candidate $C_{24}H_{21}N_3O_5S$ (rank #3 with helix) and its top stoichiometric twin.

**Query molecule properties:**

| Property | Value |
|---|---|
| Heavy atoms | 33 |
| Predicted $pIC_{50}$ (with helix) | 6.798 |
| Boltz-2 $P_{bind}$ (with helix) | 0.569 |
| Classification | Hit |
| Number of twins | 1 |

**Top-twin:**

| Twin formula | Sim. | Z | Exp. $pIC_{50}$ |
|---|---|---|---|
| $C_{39}H_{45}N_5O_8S_2$ | 0.9936 | 2.684 | 6.917 |

**Table S8.** Candidate $C_{24}H_{21}N_3O_4$ (rank #8 with helix, rank #4 without helix) and its top stoichiometric twin. The single twin achieves a perfect similarity score (1.0000) and has experimental $pIC_{50}$ = 8.482, indicating that this candidate also occupies a high-potency stoichiometric neighbourhood.

**Query molecule properties:**

| Property | Value |
|---|---|
| Heavy atoms | 31 |
| Predicted $pIC_{50}$ (with helix / without helix) | 6.241 / 6.271 |
| Boltz-2 $P_{bind}$ (with / without helix) | 0.586 / 0.571 |
| Classification | Hit (both conditions) |
| Number of twins | 1 |

**Top-twin:**

| Twin formula | Sim. | Z | Exp. $pIC_{50}$ |
|---|---|---|---|
| $C_{24}H_{29}N_3O_4$ | 1.0000 | 6.768 | 8.482 |
